# Scalp EEG detects motor sequence replay signatures during wakeful rest

**DOI:** 10.64898/2026.10.22.760953

**Authors:** Debadatta Dash, Hisato Sugata

## Abstract

Brief wakeful rest may support motor-memory processing through replay of recently practiced neural sequences, but human motor replay has been demonstrated mainly with invasive recordings or magnetoencephalography. Here we tested whether ordered motor-sequence replay can be detected from scalp EEG across wakeful rest during motor skill learning. During rest, participants received no cue, regular or accelerated auditory targeted-memory reactivation (TMR), or randomized tones. Neural states re-emerged in the practiced sequence order, revealing forward replay across conditions. Accelerated TMR enhanced total learning; within this condition, stronger forward replay showed the clearest exploratory relationship with micro-offline gains, while forward and reverse replay also covaried with gains across the cohort. The apparent replay timescale shifted systematically with decoder temporal resolution, showing that measured timing reflects both neural dynamics and the observation model. These findings support scalp EEG as a scalable approach for studying ordered motor-sequence reactivation during wakeful rest.

## Introduction

The brief pauses separating bouts of practice are increasingly recognized as an important window in motor learning. Performance can improve across rests lasting only seconds [1,2], raising the possibility that neural processes engaged after movement contribute to the evolving skill. Neural replay—the structured re-expression of experience-related neural states, often as rapidly compressed forward or reverse sequences—is a compelling candidate mechanism. Awake replay is well established in hippocampal ensembles [8,9,15]; in humans, intracortical recordings have revealed replay of learned motor-cortical firing sequences [3], while magnetoencephalography (MEG) has identified rapid hippocampo-neocortical replay during inter-practice rest that relates to motor-skill gains [4]. Together, these findings suggest that the moments between actions may contain structured neural activity relevant to how newly acquired skills are reorganized.

Yet the methods that have resolved human motor replay are difficult to deploy broadly. Intracranial recordings provide exceptional physiological specificity but are necessarily restricted to rare clinical settings, whereas MEG depends on specialized infrastructure and is challenging to scale to large cohorts. Scalp electroencephalography (EEG), by contrast, is widely available, inexpensive, portable and well suited to longitudinal and clinical studies. Task-related EEG patterns can re-emerge after learning [18], but reactivation alone is not replay: replay requires evidence that multiple neural states recur in the learned temporal order. Whether scalp EEG contains sufficient information to recover that sequential structure during waking rest therefore remains an important unresolved question. Establishing such an assay would make human replay accessible to experimental settings and populations in which invasive recordings or MEG are impractical.

Motor-sequence learning provides a particularly stringent test because the identity of an action is not sufficient to specify its role within a learned skill. In the sequence 4–1–3–2–4, for example, the same finger begins and ends the sequence, yet these two actions occupy different positions and predict different subsequent states. Recent MEG work showed that neural patterns associated with identical movements become differentiated by their sequence context during learning, with this contextualization emerging prominently across rest [11]. A replay analysis that collapses these occurrences into a single motor state may therefore discard part of the memory’s learned structure. Resolving motor replay from scalp EEG requires not only decoding the component actions but preserving their sequential context and asking whether those context-sensitive states subsequently reappear in the practiced order. Temporally delayed linear modelling (TDLM) provides a framework for quantifying such ordered transitions in decoded neural activity [5], while recent methodological work highlights the importance of understanding how decoder properties shape replay detection and its apparent timescale [6].

Targeted memory reactivation (TMR) provides an additional experimental context to examine replay and behavior. Sensory cues associated with learned material can bias subsequent memory processing, an effect studied extensively during sleep [16,17]. Wakeful TMR has recently been extended to the brief rests interleaved with motor practice, where cue timing and structure influence behavior [7]. We analyzed this dataset that comprises 64 participants practicing the same motor sequence while receiving regular, accelerated, randomized or no auditory cues during rest, together with 61-channel scalp EEG. We asked whether scalp EEG can detect learned-order reactivation during wakeful rest, whether repeated actions retain sequence-context information, how decoder temporal integration shapes the apparent replay timescale, and whether replay measures show associations with motor learning gains.

## Results

### Motor performance improves across alternating practice and rest

Participants learned the five-element sequence 4–1–3–2–4 across 36 alternating 10-s periods of practice and wakeful rest (Fig. 1a). The experiment included four conditions: no auditory cueing during rest (No cue, n = 17), replay of the learned tone sequence at the preceding practice rate (Regular, n = 16), replay of the same sequence at 1.3× the practice rate (Fast, n = 16), and randomized-pitch stimulation at regular timing (Random, n = 15; Fig. 1b). Across conditions, correct-sequence execution became progressively faster over practice, with the largest changes occurring early in learning (Fig. 1c).

**Figure 1.**
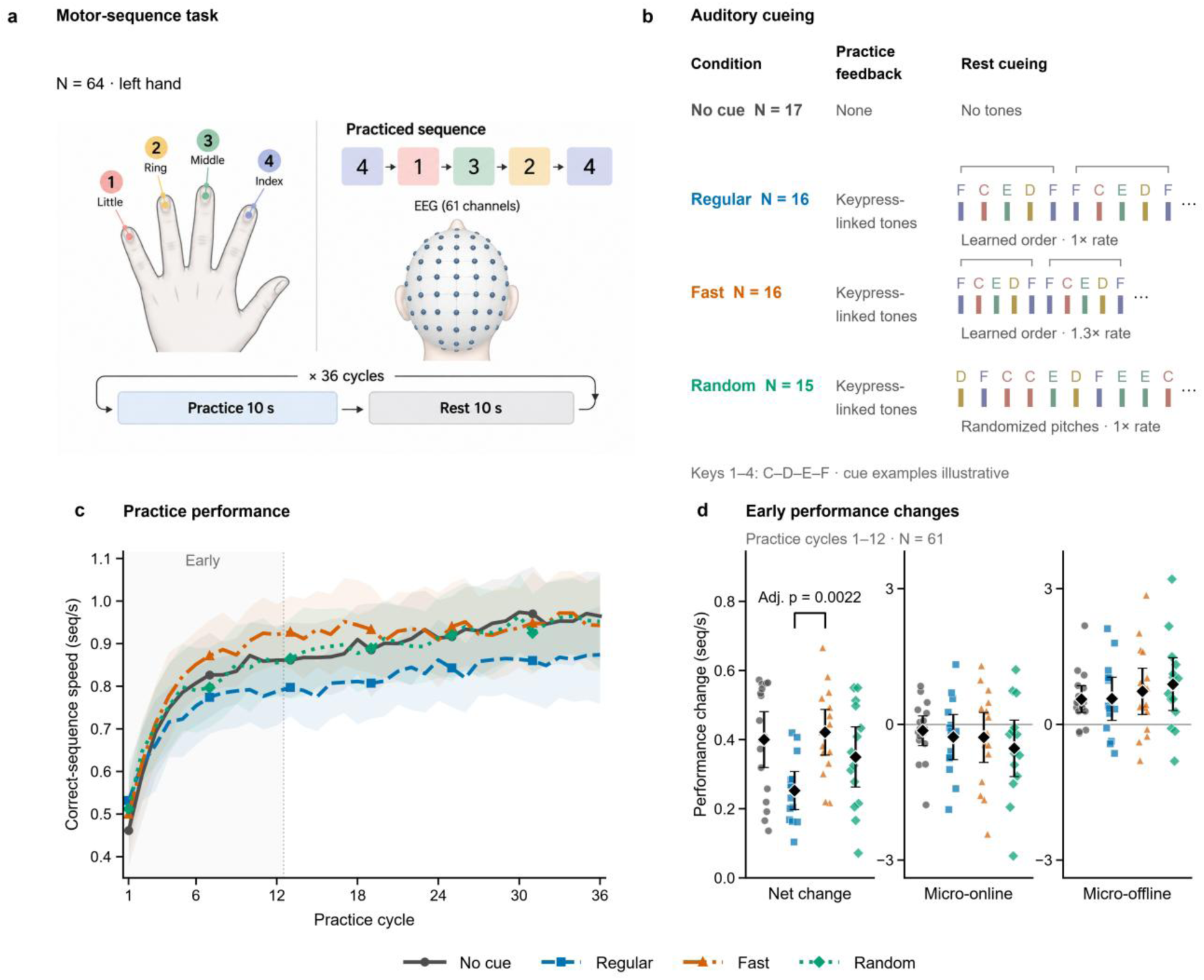
Task design, auditory cueing, and motor performance. a, Participants practiced 4–1–3–2–4 with the nondominant left hand during 36 alternating 10-s practice and 10-s rest periods while 61-channel scalp EEG was recorded. b, Auditory conditions. Regular replayed the learned tone order F–C–E–D–F at the preceding practice rate; Fast used the same order at 1.3× that rate; Random used randomized pitches at regular timing; No cue contained no rest-period auditory stimulation. c, Correct-sequence execution speed across practice. Lines show participant means and shading denotes participant-based 95% confidence intervals. d, Early performance changes over cycles 1–12. Points show participants; diamonds and whiskers denote means and 95% confidence intervals. Fast cueing produced greater early trial-mean improvement than Regular cueing (Bonferroni-adjusted p = 0.0022). MOG, micro-offline gain.

The auditory manipulation was clearly expressed early in learning. From practice cycles 1–12, participants receiving Fast cueing showed a greater increase in speed than those receiving Regular cueing (Fast–Regular difference = 0.169 sequences s⁻¹, 95% CI 0.087–0.251, Bonferroni-adjusted p = 0.0022; Fig. 1d). Early performance also showed the expected within-practice slowing and between-practice improvement: cumulative micro-online change was negative, whereas cumulative micro-offline gain was positive across groups. These behavioral measures defined the performance endpoints used in the later replay–behavior analyses. Additional behavioral reconstruction and endpoint distributions are shown in Supplementary Fig. S1.

### Scalp EEG decodes motor identity and sequence context

We next asked whether scalp EEG contained enough information about the component actions of the skill to track their re-expression during rest. Participant-specific shrinkage linear discriminant classifiers were trained on EEG from the 150 ms preceding each practice keypress, with entire practice cycles held out during cross-validation (Fig. 2a). This required the decoder to generalize to movements performed in practice cycles it had never seen.

**Figure 2.**
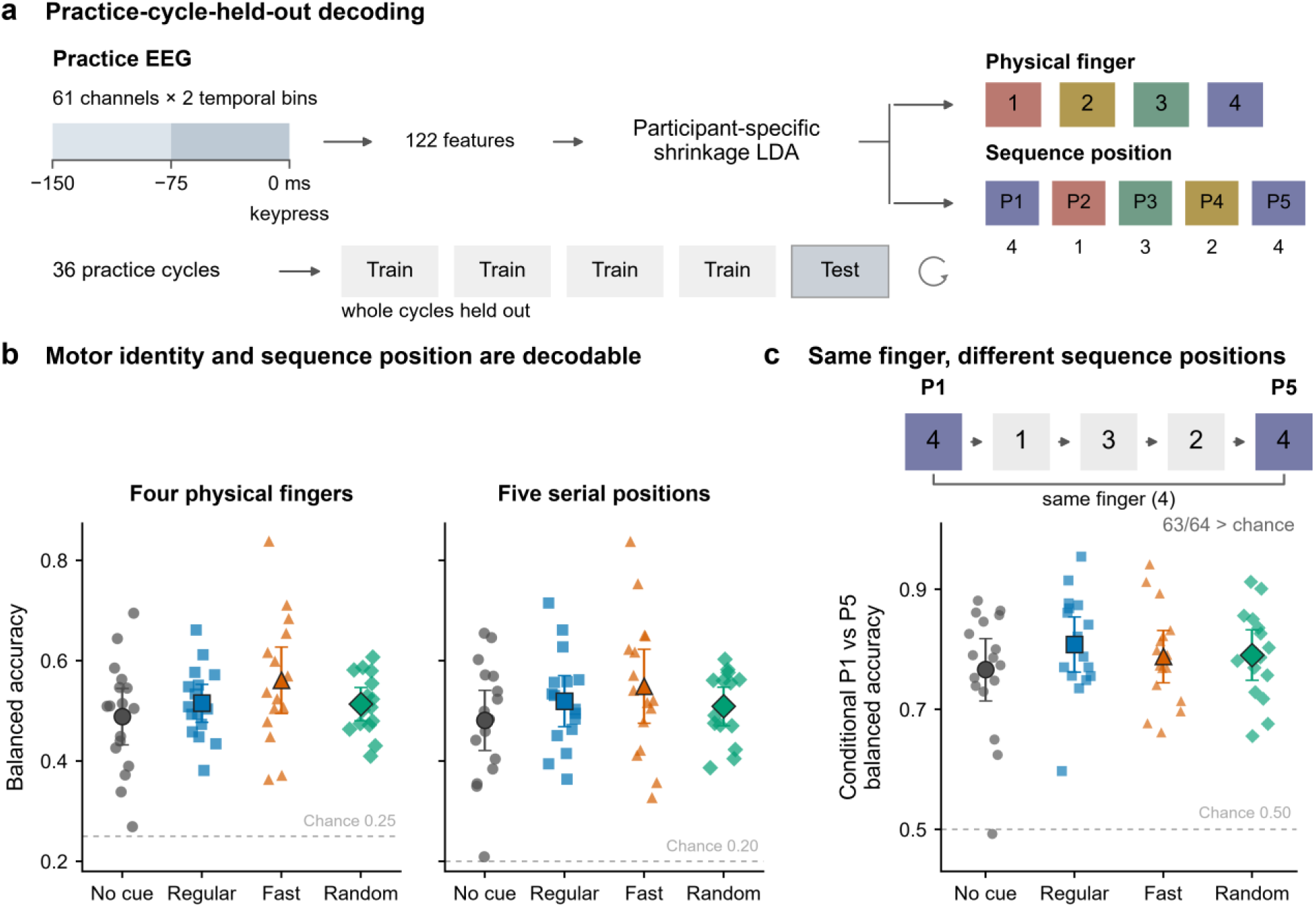
Scalp EEG distinguishes motor identity and sequence position across practice cycles. a, Participant-specific decoder framework. EEG from −150 to 0 ms before each practice keypress was summarized using two temporal-bin means across 61 scalp channels and classified with shrinkage linear discriminant analysis; complete practice cycles were held out in fivefold validation. b, Participant-level balanced accuracy for four physical fingers and five serial positions. Dashed lines indicate nominal chance (0.25 and 0.20). Small symbols denote participants; larger symbols and whiskers show group means and participant-based 95% confidence intervals. c, Conditional P1-versus-P5 discrimination from held-out five-position probabilities. P1 and P5 correspond to the same physical finger at the beginning and end of 4–1–3–2–4; nominal chance is 0.50.

Scalp EEG distinguished the four actions (physical finger state): mean balanced accuracy ranged from 0.489 to 0.561 across cueing conditions, compared with a nominal chance level of 0.25 (Fig. 2b). EEG also carried information about where an action occurred within the learned sequence. Decoding the five ordinal positions P1–P5 yielded mean balanced accuracies of 0.481–0.549 against a nominal chance level of 0.20. Observed balanced accuracy exceeded nominal chance in all 64 participants for both decoding schemes.

The repeated finger 4 provided a particularly informative test of sequence context. The first and last elements of 4–1–3–2–4 require the same physical finger but occupy different positions within the learned sequence. When held-out predictions were restricted to these two occurrences, P1-versus-P5 discrimination reached mean balanced accuracies of 0.766–0.808 across groups, and 63 of 64 participants exceeded the nominal 0.50 level (Fig. 2c). Thus, scalp EEG carried information about both movement identity and the sequential context in which that movement occurred. This allowed the complete practiced order to be represented while keeping its repeated first and final action distinct. Class-wise confusion matrices, fold stability and the P1/P5 diagnostic are shown in Supplementary Fig. S2.

### The practiced sequence re-emerges in forward order during waking rest

Having established that scalp EEG resolved both movement identity and sequence context, we asked whether the practiced transition order was detectable after movement stopped. Each participant’s practice-trained five-output decoder was applied continuously to the interleaved wakeful-rest EEG. TDLM then quantified how strongly transitions among decoded states expressed the practiced Forward sequence relative to its Reverse and an alternative Control order (Fig. 3a).

**Figure 3.**
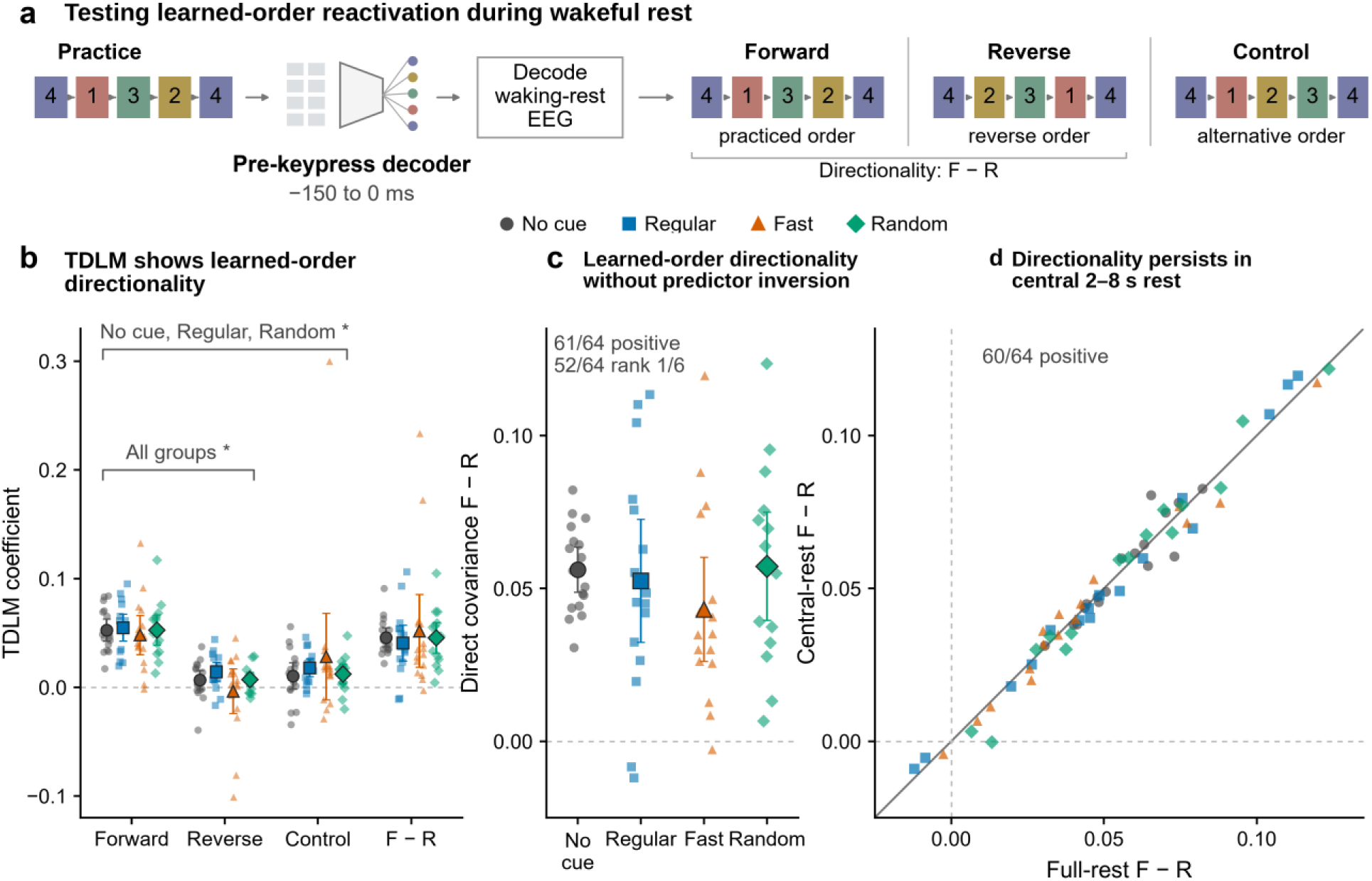
Learned-order reactivation during wakeful rest. a, Practice-trained five-output decoders were applied to wakeful-rest EEG. The representation preserves the complete learned sequence 4→1→3→2→4 while distinguishing the repeated first and final finger. Forward, Reverse and Control transition templates were evaluated at the historical 150-ms observation window and 20-ms lag. b, Participant-level TDLM Forward (F), Reverse (R), Control (C) and F−R coefficients across the four cueing groups. c, Direct lagged-covariance F−R using the same decoded EEG traces; 61 of 64 participants showed positive directionality and the practiced sequence ranked first among six constrained orders in 52 of 64. d, Participant-level full-rest versus central 2–8-s direct-covariance F−R; 60 of 64 participants retained positive directionality during central rest. Symbols denote participants and larger markers/whiskers denote group means and 95% confidence intervals where shown. Asterisks in b refer to the fixed-setting conditional learned-order reference; the reference construction and broader multiplicity sensitivities are reported in the Supplementary Information.

With a 150-ms temporal resolution decoder and 20-ms transition-lag setting, forward-minus-reverse sequenceness (F−R) was positive in every cueing condition: 0.0459 in No cue, 0.0406 in Regular, 0.0517 in Fast and 0.0453 in Random (Fig. 3b). The underlying coefficients favored Forward over Reverse, consistent with preferential expression of the practiced transition order during the brief rests separating successive bouts of practice.

The same directional organization was observed with direct lagged covariance, which measures lagged state relationships without the first-level predictor inversion used by TDLM. At 20 ms, direct-covariance F−R averaged 0.0562, 0.0525, 0.0431 and 0.0572 in No cue, Regular, Fast and Random, respectively (Fig. 3c). Sixty-one of 64 participants had a positive learned-direction contrast, and the practiced sequence ranked first among six constrained sequence orders in 52 of 64 participants. This convergence indicates that the forward learned-order signature is robust to the choice of lagged-association estimator.

Learned-order directionality was not limited to observations immediately adjacent to the practice–rest transition. Restricting the analysis to the central 2–8 s of each 10-s rest preserved the pattern: 60 of 64 participants retained positive central-rest F−R, and participant values closely followed their full-rest estimates (Fig. 3d). Together, these results identify a reproducible, directionally organized replay-related signature in scalp EEG during wakeful rest. A complementary physical-finger representation and expanded lag/estimator/rest-support analyses are shown in Supplementary Figs. S3 and S4.

### Forward full-sequence replay forms a decoder-dependent temporal ridge

We next examined the temporal organization of the replay signature. With the fixed 150-ms full-sequence decoder, Forward sequenceness rose across short lags and reached closely aligned maxima across all four experimental groups: 85 ms in No cue and Regular and 80 ms in Fast and Random (Fig. 4a). Forward-minus-Control followed a similar profile, whereas Reverse remained comparatively small. These curves reveal a coherent temporal structure in the decoded resting-state sequenceness statistic.

**Figure 4.**
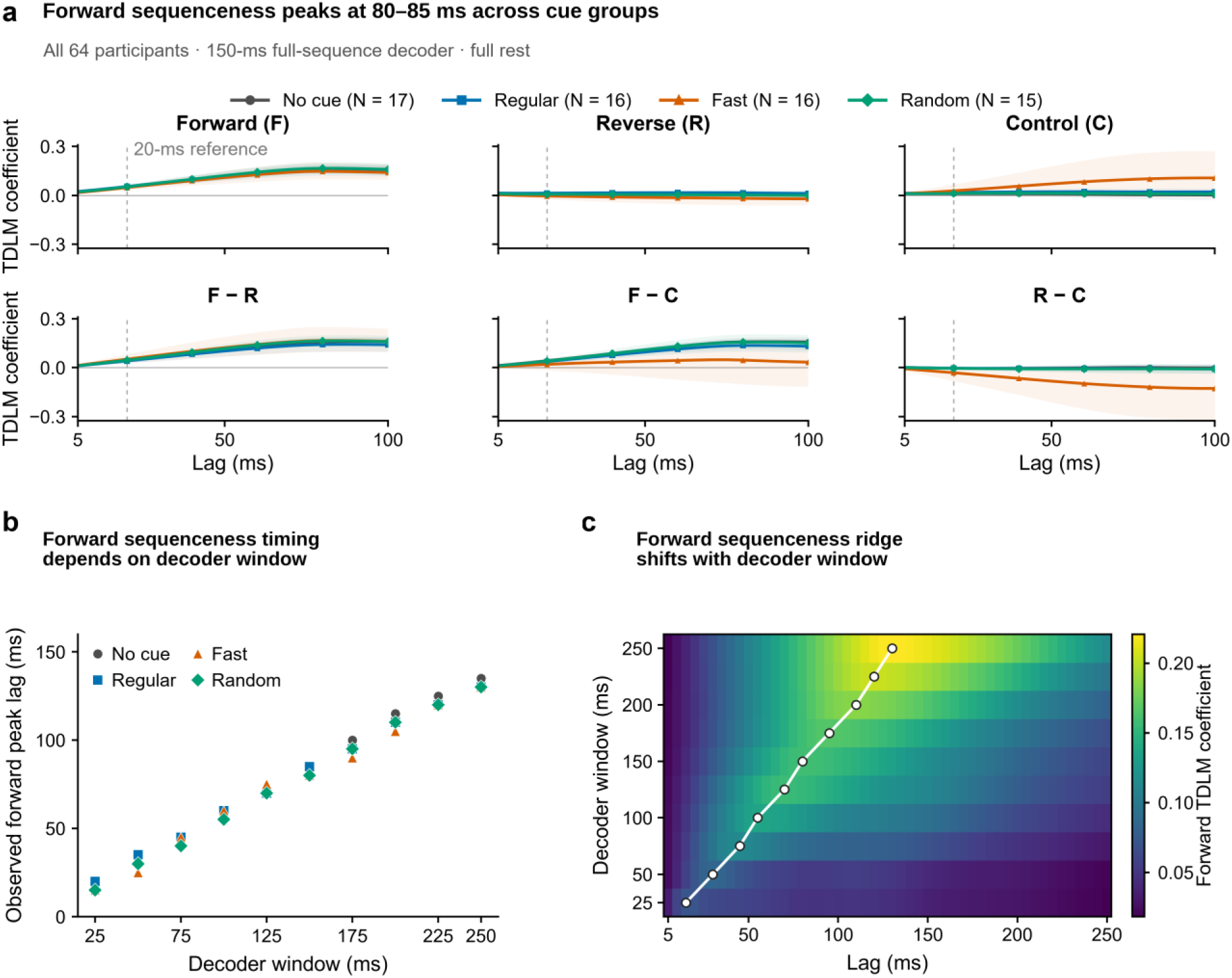
Temporal profiles of Forward sequenceness depend on decoder window. a, Lag-resolved TDLM Forward, Reverse, Control and directional contrasts from the fixed 150-ms full-sequence decoder in all 64 participants. Lines show cue-group means; shading denotes pointwise participant-bootstrap 95% confidence intervals. Dashed lines mark the historical 20-ms lag. Forward maxima occurred at 80–85 ms across groups. b, Group-specific Forward peak lag as a function of decoder observation window from 25 to 250 ms. c, Pooled Forward TDLM window × lag surface across all 64 participants. White markers denote each window’s observed Forward maximum, which shifted from 15 ms at W25 to 130 ms at W250; the pooled W150 maximum was 80 ms. Peak lag is interpreted as a property of the decoded sequenceness statistic rather than an intrinsic replay speed.

A broader analysis across decoder observation windows showed that this temporal maximum moved systematically with decoder duration. Across all 64 participants, short-window decoders produced early Forward maxima, whereas progressively longer windows shifted the maximum to later lags (Fig. 4b). In the pooled window × lag surface, the Forward peak moved from 15 ms with a 25-ms decoder to 130 ms with a 250-ms decoder, passing through 80 ms at the fixed 150-ms decoder used in the primary analysis (Fig. 4c).

Rather than producing isolated peaks, the data formed a continuous temporal ridge across decoder resolutions. This relationship provides an important measurement-level insight: the temporal structure detected as replay is strongly coupled to the temporal integration of the neural-state decoder itself. Scalp EEG therefore reveals a robust sequential signal during rest, while the timescale at which that signal is expressed must be interpreted in the context of the decoder used to observe it. Decoder-window decomposition and decoded-state autocorrelation are shown in Supplementary Fig. S6.

### Forward and reverse replay moderately correlate with motor learning gains across lags

We next explored whether individual differences in sequenceness during rest were associated with the behavioral gains that emerged across the same rest periods. Micro-offline gain was defined as the change from the final correctly executed sequence before a rest to the first correctly executed sequence in the following practice period. Early cumulative gain summed the first 11 inter-practice transitions, whereas total cumulative gain summed all 35. Complete behavioral endpoints were available for 61 participants.

Across the full cohort, both Forward and Reverse replay showed moderate positive relationships with motor gains over neighboring transition lags rather than at isolated time points (Fig. 5a). For early gain, the largest ordinary associations occurred at 30 ms for Forward (Spearman ρ = 0.283, p_nom = 0.032) and 35 ms for Reverse (ρ = 0.307, p_nom = 0.018). After adjustment for cueing condition and initial performance, the early Reverse relationship reached partial ρ = 0.315 at 65 ms (p_nom = 0.020). These were nominal associations; The relationships didn’t survive under strict Family Wise Error Rate (FWER160) correction. We therefore treat the lag-resolved replay–behavior analysis as exploratory.

**Figure 5.**
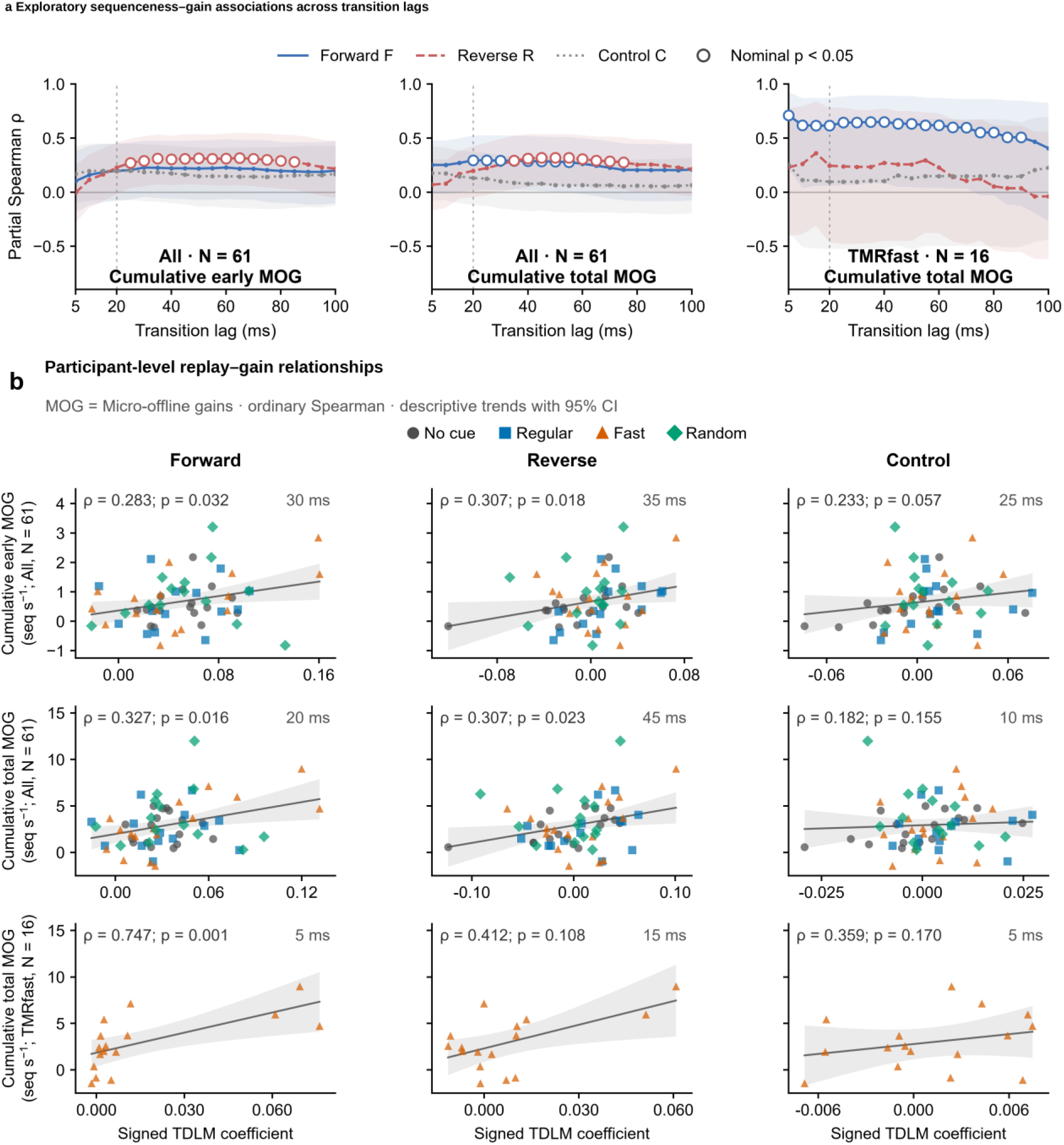
Exploratory sequenceness–motor gain associations. a, Lag-resolved partial Spearman associations between four-physical-finger TDLM Forward (F), Reverse (R) and Control (C) coefficients and reconstructed cumulative micro-offline gain. Left, early gain across rests 1–11 in all participants with complete endpoints (N = 61); center, total gain across rests 1–35 in the same cohort; right, total gain within Fast cueing (N = 16). Curves are adjusted for baseline performance and, for pooled analyses, cueing condition; shading denotes pointwise participant-bootstrap 95% confidence intervals and open circles mark nominal permutation p < 0.05. b, Participant-level relationships at selected ordinary Spearman lags; points are participants and lines are descriptive fits. The Fast-group Forward association at 5 ms (ρ = 0.747, p_nom = 0.001, *p_Holm = 0.032, p_FWER960 = 0.165*) is an exploratory subgroup result and did not survive the FWER960 reference (p = 0.165). MOG, micro-offline gain.

The relationship was also evident when gains were accumulated across the full session. Adjusted Forward replay was positively related to total cumulative gain over lags from 20–55 ms and reached partial ρ = 0.295 at 20 ms (p_nom = 0.031). Reverse replay was similarly related to total gain over 35–75 ms, reaching partial ρ = 0.320 at 50 ms (p_nom = 0.018). This association also did not survive FWER160 correction. Participant-level scatter plots show the underlying distributions at retrospectively selected lags with descriptive visualizations of the nominal positive association (Fig. 5b). The corresponding full-sequence associations are shown in Supplementary Fig. S7.

The strongest participant-level replay–gain relationship emerged in the Fast-cueing condition, which also showed the clearest early behavioral advantage in Fig. 1. Among the 16 Fast participants, Forward replay correlated positively with total cumulative micro-offline gain (ρ = 0.747 at 5 ms, nominal permutation p = 0.0008; Fig. 5a,b). The association remained positive across neighboring lags and was descriptively stronger than the corresponding Reverse and Control relationships. It survived Holm correction across 20 lags and two outcomes: early and total (40 tests; p_Holm = 0.032), but not the broader correction spanning all four groups, both adjustment models, three components and two outcomes (FWER960 p = 0.165). Together, the behavioral analyses provide a functional bridge between the neural sequence dynamics observed during waking rest and the performance improvements that accumulate across practice, with the clearest exploratory coupling emerging for Forward replay in the Fast-cueing group.

## Discussion

### Scalp EEG detects sequence-level replay signatures during wakeful rest

The central advance of this study is that learned-order reactivation of a recently practiced motor sequence can be detected from scalp EEG during the seconds between bouts of practice. Human motor replay has previously been demonstrated with intracortical recordings during rest and with MEG during motor learning, while non-spatial MEG studies have shown that internally generated state sequences can unfold rapidly in humans [3,4,10]. TDLM provides a general framework for quantifying such ordered reactivation from decoded neural states [5]. Our findings extend this literature by showing that learned order replay signature is accessible with conventional scalp electrophysiology in a comparatively large cohort and during the brief wakeful rests naturally embedded within skill acquisition.

Replay is more than recurrence of individual states: its defining feature is temporal organization. Awake hippocampal replay can preserve the order of recent experience in forward and reverse directions, and sequence order has been proposed to support computations over relationships among states rather than simple reinstatement of isolated representations [8,9,15]. In the present motor task, decoded states expressed the transition structure of 4–1–3–2–4 after overt movement stopped. The directional signature was present during uncued rest and across the three auditory-cueing conditions and remained detectable within the central portion of the rest interval. Together, these observations indicate that short pauses in practice contain structured neural dynamics that retain information about the recently expressed action sequence.

The ability to measure this organization with scalp EEG broadens the experimental reach of human replay research. Intracranial recordings and MEG provide high-value mechanistic and temporal information but constrain accessibility and repeated testing. A scalp-EEG assay of ordered reactivation could support larger cohorts, longitudinal measurements and studies of populations in whom invasive or MEG recordings are impractical. The present work therefore contributes both a biological observation and a scalable measurement framework for studying sequence-resolved neural dynamics during waking motor memory processing.

### Sequence context is part of the representation that re-emerges during rest

A particularly informative feature of the learned sequence is that finger 4 occurs at both its beginning and its end. The physical action is identical, yet EEG distinguished these two occurrences across held-out practice cycles. This finding converges with prior MEG evidence that the neural representation of the same action differentiates according to its location within a learned sequence [11]. Motor-state representations therefore carry information about sequential context in addition to movement identity.

Contextualization changes how replay should be interpreted. Sequence learning embeds individual actions within a larger temporal and predictive structure, so the same movement can acquire different neural context depending on what preceded it and what is expected to follow. Human MEG studies have similarly shown rapid sequences of non-spatial state representations during periods without external stimulation [10], while hippocampal work demonstrates internally generated trajectories directed toward remembered future goals [12]. When context-sensitive motor states subsequently recur in their practised order, the resulting signal is therefore more naturally viewed as re-expression of an organized action sequence than as independent recurrence of isolated motor commands.

This observation also defines a clear next experimental step. Counterbalancing the learned sequence while preserving the same component movements would separate memory-specific transition structure from fixed task context, and independent movement localizers would further dissociate action identity from ordinal context. Such designs should allow future studies to ask which level of the motor representation—individual movement, ordinal context, transition structure or an integrated sequence—is preferentially reinstated during waking rest.

### Replay timing reflects both neural dynamics and the observation model

A second conceptual contribution concerns the temporal interpretation of replay. With the fixed 150-ms decoder, Forward sequenceness showed a reproducible maximum around 80–85 ms across cueing conditions. In many replay studies, short transition times are biologically informative because internally generated sequences unfold faster than overt behavior [8–10,15]. Taken in isolation, the present maximum could therefore be tempting to interpret as an intrinsic transition time or compression rate.

The decoder-window analysis shows why that inference would be premature. As the temporal observation window used to define a neural state increased, the apparent Forward maximum shifted systematically to longer lags. The measured peak thus formed a continuous ridge whose location depended partly on the temporal integration of the decoder. This result is consistent with the general principle that TDLM operates on classifier-derived state evidence [5] and with recent hybrid-simulation work showing that replay detectability and inferred sequenceness depend strongly on the properties of the decoding and analysis pipeline [6]. The temporal properties of the measurement can therefore become embedded in the temporal properties attributed to replay.

Decoder dependence does not remove the evidence for ordered reactivation. A coherent Forward ridge persisted across a broad range of decoder resolutions; what changed was the precise lag at which the statistic was maximal. The defensible conclusion is therefore that recently practiced motor states exhibit reliable short-lag forward temporal organization during wakeful rest, rather than that replay occurs at one privileged 80-ms interval. Comparisons of replay timing across studies, recording modalities or decoder architectures should explicitly account for the temporal resolution of the state representation used to measure it.

### Forward and reverse replay may support complementary computations during learning

The relationship between replay and behavior provides an important indication that these resting dynamics may be functionally relevant. Across participants, Forward and Reverse sequenceness was positively associated with cumulative micro-offline motor gains over neighboring transition lags. This broadly echoes the earlier MEG finding that waking motor replay covaries with early skill acquisition [4], while extending the association to scalp EEG and to continuous lag-resolved sequence measures. However, none of the lag-specific associations survived a stricter family-wise correction in the present dataset. The behavioral results therefore warrant caution to establish a robust replay–behavior relationship.

The involvement of both replay directions is biologically interesting rather than contradictory. Forward and reverse replay are well established in hippocampal circuits [8,9]. Their functional profiles are not identical: forward hippocampal sequences can represent upcoming goal-directed paths [12], whereas reverse replay is particularly sensitive to changes in reward value [13]. Computational accounts likewise suggest that forward and reverse replay can emerge from different priorities for prospective planning and backward propagation of newly acquired information [14]. These findings do not imply that motor replay serves the same functions as spatial hippocampal replay, but they provide a principled reason not to treat Reverse replay simply as noise or the absence of Forward replay.

In a motor-learning setting, the two directions could therefore reflect complementary operations on the developing sequence - for example, prospective organization of upcoming actions and retrospective updating of recently executed transitions. The present data cannot distinguish these computations, but the positive behavioral relationships motivate experiments that manipulate sequence direction or task demands prospectively. Importantly, the behavioral endpoint here is micro-offline gain. Brief-rest gains were originally interpreted as rapid offline consolidation [1], whereas recent debates indicate that they can also partly reflect transient performance recovery and advance motor planning rather than durable sequence-specific learning [2]. We therefore interpret the present associations as links between replay-related dynamics and short-timescale motor skill improvement, not as direct evidence that replay produced lasting consolidation.

### Accelerated memory reactivation and replay-behavior coupling

The Fast cueing condition provides an especially intriguing extension of this functional relationship. In the parent experiment, replaying the learned auditory sequence faster than the preceding practice rate enhanced overall motor learning relative to regular-rate cueing [7]. In the present analysis, the same condition showed moderate yet nominally significant positive participant-level relationship between Forward replay and cumulative motor gain. Notably, this pattern emerged in replay-behavior coupling rather than as a simple increase in the group-average directional replay statistic. This distinction suggests a useful biological hypothesis: targeted reactivation may influence learning not only by changing how much sequence activity occurs, but by changing how effectively that activity is coupled to the evolving memory trace. Most targeted-memory-reactivation evidence comes from sleep, where behavioral effects vary with memory domain, sensory cueing protocol and physiological state [16,17]. The wakeful motor paradigm studied here is therefore a distinct setting in which cue rate can be related directly to ongoing practice and to replay measured in the immediately intervening rest [7]. Faster cueing could alter the temporal alignment between external stimulation, endogenous sequence dynamics and the current state of the motor memory, making replay more behaviorally consequential without necessarily increasing its overall magnitude.

This interpretation is a testable hypothesis rather than a demonstrated mechanism. A prospective within-participant design could manipulate cue rate, preserve precise cue timestamps and quantify replay relative to each participant’s current sequence speed. If replay-behavior coupling varies systematically with the temporal relationship between cueing and endogenous sequence dynamics, it would provide a mechanistic account of when wakeful TMR facilitates learning and why its effects differ across individuals.

### A scalable framework for human motor-memory replay

More broadly, the present findings position scalp EEG as a practical platform for studying human replay beyond specialized recording environments. The combination of practice-trained decoding, sequence-context representation and lagged transition analysis can be repeated across learning, across days and across populations. This creates opportunities to examine replay during neurorehabilitation, developmental learning and longitudinal skill acquisition, and to test whether behavioral or neuromodulatory interventions alter the relationship between replay and subsequent performance.

The study also suggests several methodological principles for that broader use. Replay should be evaluated at the level of sequence organization rather than inferred from isolated increases in decoded state evidence; Forward, Reverse and alternative transition structures should be examined separately; and apparent replay timescales should be interpreted in light of decoder temporal resolution. These principles make the assay more conservative, but they also make the resulting biology more interpretable.

Several features bound the interpretation. The primary measure is continuous learned-order sequenceness rather than independently verified discrete replay events. Nevertheless, the forward sequence preference was reproduced with both TDLM and direct lagged covariance, was positive in most participants, and persisted in the central portion of rest, supporting a robust replay-related sequence signature in scalp EEG. Because all participants learned the same 4–1–3–2–4 sequence, memory-specific content cannot be completely separated from stable task structure. Counterbalanced learned sequences and independent state localizers will be important for establishing stronger content specificity.

The exact temporal scale of replay also remains method dependent. Forward sequenceness showed a coherent temporal ridge, but its peak shifted systematically with decoder observation window; therefore, the observed lag maximum should not be interpreted as an intrinsic physiological replay speed.

Finally, the replay–behavior relationships are exploratory and observational. Forward and reverse sequenceness covaried with micro-offline motor gains but did not survive corrections. Positive replay-behavior association in the Fast TMR condition is obtained with limited sample size (N=16). Thus, these findings do not establish that replay causes learning or that accelerated cueing acts through replay. Prospective manipulation of replay and delayed retention testing will be needed to establish causality.

Taken together, the results identify a reproducible signature of ordered motor-sequence reactivation in human scalp EEG and connect that signature to the behavioral dynamics of skill acquisition. The broader implication is that the short rests interleaved with practice are crucial for learning: they contain structured re-expression of the recently practiced sequence, and the strength and direction of that organization carry information about how performance changes across learning. By making these dynamics accessible with scalp EEG, this work provides a foundation for larger-scale and ultimately mechanistic studies of human motor-memory replay.

## Methods

### Participants and experimental design

We reanalyzed scalp EEG from the wakeful targeted-memory-reactivation motor-learning study described previously [7]. The dataset comprised 64 participants assigned to four rest conditions: No cue (n = 17), Regular auditory cueing (n = 16), Fast auditory cueing (n = 16), and Random cueing (n = 15). Participants practiced the left-hand sequence 4–1–3–2–4 across 36 cycles, each consisting of 10 s of practice followed by 10 s of waking rest. During rest, Regular cueing replayed the learned auditory sequence at the preceding practice rate, whereas Fast cueing replayed the same sequence at 1.3× that rate. Random cueing presented randomized tones at the regular timing, and No cue contained no auditory stimulation. Recruitment, consent, ethical approval and the original experimental protocol are described in the parent study [7]. All available participants and all 36 rest periods were retained for the neural analyses; no participant was excluded on the basis of decoder performance or replay-effect direction.

### EEG preprocessing

Recordings contained 61 scalp EEG channels together with auxiliary physiological channels. Continuous EEG was band-pass filtered from 0.5–100 Hz, notch filtered at 60 Hz, processed using independent-component analysis with automated component classification, and represented at 600 Hz for the present analyses. Only the 61 scalp EEG channels entered the neural decoders. Practice keypresses and rest boundaries were aligned using the task markers supplied with the recordings. Preprocessing was performed on the continuous recordings before neural-state decoding. Detailed preprocessing parameters, component-processing procedures, auxiliary-channel analyses and quality-control results are provided in Supplementary Methods.

### Behavioral measures

Correct instances of the instructed 4–1–3–2–4 motif were identified from recorded keypresses. Sequence speed was defined as the reciprocal of the elapsed time between the first and fifth presses of a correctly completed motif and expressed in sequences s⁻¹. Practice-cycle performance was summarized as the mean speed of correctly completed motifs within each cycle. For the early behavioral analysis, trial-mean learning was quantified as the difference between mean sequence speed in practice cycle 12 and cycle 1. Micro-online change within each practice period was defined as the speed of the last correct motif minus that of the first correct motif. Micro-offline gain (MOG) across rest i was defined as the first correct sequence speed in practice i+1 minus the last correct sequence speed in practice i. Early cumulative MOG summed the first 11 practice-to-practice transitions, spanning cycles 1–12. Total cumulative MOG summed all 35 transitions across the session. Early cumulative micro-online change was obtained by summing within-practice changes across cycles 1–12. Three participants—two in Regular and one in Random—did not have the complete cycle-1 correct-motif information required for the reconstructed early and session behavioral endpoints. Missing values were not imputed, yielding 61 participants for the replay–behavior analyses (No cue, n = 17; Regular, n = 14; Fast, n = 16; Random, n = 14). The original behavioral scoring code from the parent study was not part of the delivered dataset; validation of the reconstructed measures against reported behavioral summaries and additional behavioral definitions are provided in Supplementary Methods. Between-group comparisons of the reconstructed phase-specific trial-mean learning measures used two-sided Welch tests. The nine auditory-group pair × learning-phase comparisons were Bonferroni corrected as a single family.

### Neural-state feature extraction and temporal alignment

Participant-specific neural-state decoders were trained from EEG immediately preceding practice keypresses. For each labeled movement, the primary decoder used the 150-ms interval preceding the keypress (−150 to 0 ms), selected from a validation controlled replay injection experiment across 10 temporal resolutions (25 to 250ms with 25 ms increments) and three alignments (Pre-onset, peri-onset, and post-onset). This interval was divided into two consecutive 75-ms bins, and mean voltage was calculated separately within each bin for each of the 61 EEG channels. Each movement was therefore represented by 122 features. The two-bin representation retained coarse temporal information within the pre-keypress period while keeping feature dimensionality modest. We defined two complementary state spaces. The four-class decoder represented physical finger identity (1–4). The primary full-sequence decoder represented the five ordinal elements P1–P5 of the practiced sequence, corresponding to 4–1–3–2–4. P1 and P5 were therefore assigned separate neural-state labels despite requiring the same physical finger. This allowed the full-sequence representation to retain information about where the repeated action occurred within the learned sequence. The identical feature construction was used when applying the trained models to waking-rest EEG. At each analysis time point, the decoder received the immediately preceding observation window, and probability estimates were generated every 5 ms. Observation windows were required to remain within the relevant rest interval.

### Decoder training and cross-validation

Features were standardized using parameters estimated from the training data only. Participant-specific shrinkage linear discriminant analysis classifiers were then fitted using class-balanced training samples. Decoder generalization was evaluated with fivefold cross-validation in which entire practice cycles were allocated to mutually exclusive folds. All movement events from a held-out cycle were therefore excluded from classifier training, rather than randomly mixing temporally neighboring events from the same practice period across training and test sets. Balanced accuracy was calculated from the out-of-fold predictions separately for the four-finger and five-position representations. Because the first and final elements of 4–1–3–2–4 involve the same physical finger, we additionally quantified whether the five-output decoder distinguished P1 from P5. This diagnostic used the held-out posterior probabilities for the two relevant classes and evaluated the conditional probability P5/(P1+P5) at a threshold of 0.5; no separate binary classifier was fitted. After cross-validation, participant-specific decoders were fitted to the available practice data using the same preprocessing and feature definition and then applied continuously to waking-rest EEG. Rest data were not used for classifier training or model selection.

### Full-sequence replay analysis

Replay was quantified from the decoded resting-state probability time series using temporally delayed linear modelling (TDLM) [5]. Within each rest period, decoder probabilities were normalized and standardized by state. At each tested transition lag, the lagged state-probability vector was regressed simultaneously on the concurrent state probabilities. The resulting lagged association matrix was then projected jointly onto templates representing Forward, Reverse and Control transitions, together with self-transition and general-transition terms. For the five-output full-sequence representation, the practiced Forward template was P1→P2→P3→P4→P5, corresponding to 4→1→3→2→4. The Reverse template was P5→P4→P3→P2→P1, and the Control sequence was P1→P2→P4→P3→P5. The corresponding physical-finger templates were 4→1→3→2→4, 4→2→3→1→4 and 4→1→2→3→4, respectively. Forward (F), Reverse (R) and Control (C) coefficients were retained separately, together with the directional contrasts F−R, F−C and R−C. The primary directionality measure was F−R. The historical reference analysis used the 150-ms decoder and a 20-ms transition lag. This setting was retained for the fixed-setting comparisons across the four cueing conditions, while broader lag-resolved analyses were used to characterize the temporal profile of the replay statistic.

### Direct lagged covariance

We used direct lagged covariance as a complementary measure of learned-order directionality. For each rest and transition lag, source and target state-probability vectors were centered and their cross-state covariance calculated directly. Forward and Reverse scores were obtained by averaging covariance values over the corresponding prespecified transition edges, and their difference yielded the direct-covariance F−R contrast. Unlike the first-level TDLM regression, this calculation does not require inversion of the covariance among predictor probabilities. It therefore provides a complementary estimate of directional sequence organization from the same decoded resting activity. Raw coefficient magnitudes from TDLM and direct covariance were not compared because the two estimators have different numerical scales. Participant-level scores were obtained by averaging across the available rest periods, with participants—not rest intervals or EEG samples— treated as the independent sampling units.

### Learned-order reference

To evaluate whether the practiced sequence was preferentially expressed relative to alternative state orders, we compared the observed template with six constrained sequence assignments. These alternatives preserved the sequence endpoints while permuting the intermediate state identities, thereby retaining the same decoded states but changing their order. Reference distributions were generated from 5,000 participant-wise assignments, with each participant’s state mapping retained consistently across the comparisons. Group-level statistics for the practiced order were evaluated against the corresponding maximum-statistic reference distribution. Detailed construction of the template family and extended multiplicity sensitivities are described in Supplementary Methods.

### Lag-resolved replay and decoder temporal resolution

We first characterized replay over transition lags from 5 to 100 ms in 5-ms increments using the fixed 150-ms decoder. Forward, Reverse, Control, F−R, F−C and R−C were retained across the complete lag grid. To determine how the measured temporal profile depended on the temporal resolution of the neural-state decoder, we repeated the decoding and replay analysis using observation windows from 25 to 250 ms in 25-ms increments. Each decoder window immediately preceded the practice keypress and was divided into two chronological temporal bins using the same feature construction as the primary decoder. Participant-specific models were trained using the same practice-cycle-held-out procedure and applied to rest on the same 5-ms grid. For the decoder-resolution analysis, replay was evaluated over transition lags from 5 to 250 ms. For each decoder window, the observed Forward peak was defined as the lag yielding the largest group-mean Forward coefficient. The resulting window × lag surface was used to assess how the apparent temporal structure of replay changed with decoder temporal integration. Peak lags were treated as properties of the measured replay statistic rather than assumed physiological replay durations. Additional analyses of F−R decomposition, decoded-state autocorrelation and physical-finger representation sensitivity are provided in the Supplementary Information.

### Rest-boundary and movement sensitivity analyses

As a temporal sensitivity analysis, replay was also evaluated within the central 2–8 s of each 10-s rest period, requiring the complete decoder observation window and both lagged states to remain within the same retained interval. Additional analyses censored recorded rest-period keypresses and used available hand-EMG signals to assess sensitivity to overt movement-related activity. These analyses and their availability criteria are described in Supplementary Methods. The paired keypress- and EMG-based sensitivities are shown in Supplementary Fig. S5.

### Exploratory sequenceness–behavior relationships

We tested whether individual differences in resting sequenceness were associated with cumulative MOG. Neural predictors were averaged over the same rest periods contributing to each behavioral outcome: rests 1–11 for early MOG and rests 1–35 for total MOG. Behavioral associations were evaluated for both decoder representations. The four-finger TDLM results are shown in the main Figure 5, while the full-sequence behavioral analyses are shown in Supplementary Fig. S7. Forward, Reverse, Control and F−C coefficients were examined over transition lags from 5 to 100 ms. Ordinary associations were quantified using Spearman rank correlations. Adjusted analyses used partial Spearman correlations obtained by ranking the neural predictor, behavioral outcome and baseline speed and residualizing the predictor and outcome against auditory-condition indicators and ranked initial sequence speed. Within the Fast subgroup, the corresponding adjusted analysis controlled for initial speed. Nominal significance for pooled ordinary correlations was evaluated using group-preserving permutations of the behavioral outcome. Adjusted analyses used within-group Freedman–Lane residual permutations. Each test used 5,000 resampled assignments together with the observed assignment. Pointwise 95% confidence intervals were obtained from 2,000 participant-bootstrap resamples stratified by auditory condition. The main lag-profile analyses examined four predictors (F, R, C and F−C), 20 transition lags and two behavioral outcomes, yielding 160 comparisons within each decoder representation and adjustment model. Family-wise inference used the maximum statistic across this set; extended cross-representation and group-specific sensitivity analyses are reported in Supplementary Figs. S7 and S8. Participant-level scatter plots were generated from the actual paired neural and behavioral measurements at retrospectively selected lags corresponding to the largest positive ordinary Spearman association for the displayed measure and outcome. These scatters were used to visualize the underlying participant distributions rather than as independent confirmatory tests. Group-specific relationships, including the Fast-cueing result shown in Figure 5, were treated as exploratory subgroup analyses and were not interpreted as formal between-condition interactions.

### Statistical analysis

Participants were the independent sampling units for all population-level analyses. Group summaries were calculated from participant-level estimates. Unless otherwise stated, uncertainty was expressed as participant-based 95% confidence intervals; bootstrap-based intervals used 2,000 participant resamples. All statistical tests were two-sided. Full tested lag grids were retained, including boundary maxima and negative coefficients, and no participant, lag or neural measure was excluded because of the magnitude or direction of an observed effect. Nominal and multiplicity-adjusted probabilities are distinguished explicitly throughout the manuscript. The primary replay analyses used the five-output full-sequence representation; corresponding four-finger analyses served as representation sensitivities except for the exploratory replay–behavior analysis shown in Figure 5. Detailed preprocessing specifications, permutation families and movement sensitivities are provided in the Supplementary Information; the operational candidate-event analysis is shown in Supplementary Fig. S9.

## Data Availability

Behavioral and EEG data are available from the corresponding author, Hisato Sugata, upon reasonable request.

## Code Availability

Custom analysis and figure-generation code will be made publicly available in a versioned GitHub repository upon publication.

## Author Contributions

Debadatta Dash: Conceptualization, Methodology, Software, Formal analysis, Visualization, Writing – original draft, Writing – review & editing. Hisato Sugata: Experimental design, Data collection, Writing – review & editing.

## Competing Interests

The authors declare no competing interests.

## Notes

### Competing Interest Statement

The authors have declared no competing interest.

## References

1. Bönstrup, M. et al. A rapid form of offline consolidation in skill learning. Current Biology 29, 1346–1351.e4 (2019). 10.1016/j.cub.2019.02.049

2. Das, A., Karagiorgis, A., Diedrichsen, J., Stenner, M.-P. & Azañón, E. Micro-offline gains do not reflect offline learning during early motor skill acquisition in humans. Proceedings of the National Academy of Sciences USA 122, e2509233122 (2025). 10.1073/pnas.2509233122

3. Eichenlaub, J.-B. et al. Replay of learned neural firing sequences during rest in human motor cortex. Cell Reports 31, 107581 (2020). 10.1016/j.celrep.2020.107581

4. Buch, E. R., Claudino, L., Quentin, R., Bönstrup, M. & Cohen, L. G. Consolidation of human skill linked to waking hippocampo-neocortical replay. Cell Reports 35, 109193 (2021). 10.1016/j.celrep.2021.109193

5. Liu, Y. et al. Temporally delayed linear modelling (TDLM) measures replay in both animals and humans. eLife 10, e66917 (2021). 10.7554/eLife.66917

6. Kern, S. et al. Challenges in replay detection by TDLM in post-encoding resting state. eLife 14, RP108023 (2026). 10.7554/eLife.108023

7. Kawasoe, R. et al. Wakeful targeted memory reactivation during short rest periods modulates early motor learning. npj Science of Learning 11, 23 (2026). 10.1038/s41539-026-00407-9

8. Foster, D. J. & Wilson, M. A. Reverse replay of behavioural sequences in hippocampal place cells during the awake state. Nature 440, 680–683 (2006). 10.1038/nature04587

9. Diba, K. & Buzsáki, G. Forward and reverse hippocampal place-cell sequences during ripples. Nature Neuroscience 10, 1241–1242 (2007). 10.1038/nn1961

10. Kurth-Nelson, Z., Economides, M., Dolan, R. J. & Dayan, P. Fast sequences of non-spatial state representations in humans. Neuron 91, 194–204 (2016). 10.1016/j.neuron.2016.05.028

11. Dash, D. et al. Sequence action representations contextualize during early skill learning. eLife 13, RP102475 (2025). 10.7554/eLife.102475

12. Pfeiffer, B. E. & Foster, D. J. Hippocampal place-cell sequences depict future paths to remembered goals. Nature 497, 74–79 (2013). 10.1038/nature12112

13. Ambrose, R. E., Pfeiffer, B. E. & Foster, D. J. Reverse replay of hippocampal place cells is uniquely modulated by changing reward. Neuron 91, 1124–1136 (2016). 10.1016/j.neuron.2016.07.047

14. Mattar, M. G. & Daw, N. D. Prioritized memory access explains planning and hippocampal replay. Nature Neuroscience 21, 1609–1617 (2018). 10.1038/s41593-018-0232-z

15. Foster, D. J. Replay comes of age. Annual Review of Neuroscience 40, 581–602 (2017). 10.1146/annurev-neuro-072116-031538

16. Cellini, N. & Capuozzo, A. Shaping memory consolidation via targeted memory reactivation during sleep. Annals of the New York Academy of Sciences 1426, 52–71 (2018). 10.1111/nyas.13855

17. Hu, X., Cheng, L. Y., Chiu, M. H. & Paller, K. A. Promoting memory consolidation during sleep: a meta-analysis of targeted memory reactivation. Psychological Bulletin 146, 218–244 (2020). 10.1037/bul0000223

18. Murphy, M., Stickgold, R., Parr, M. E., Callahan, C. & Wamsley, E. J. Recurrence of task-related electroencephalographic activity during post-training quiet rest and sleep. Scientific Reports 8, 5398 (2018). 10.1038/s41598-018-23590-1

