## supplementary materials for "Scalp EEG detects motor sequence replay signatures during wakeful rest"

This Supplementary Information accompanies the main manuscript with analyses that establish the behavioral and decoding foundations of the replay assay, test stability across state representations and lagged-association estimators, characterize decoder-dependent temporal resolution, and provide the complete exploratory replay–behavior analyses. Supplementary Figures S1–S9 are supported by six summary tables and machine-readable source data. Supplementary references are numbered independently.

The analyses follow the same scientific progression as the paper: behavioral and decoding foundations (Supplementary Figs. S1–S2), representation and temporal structure of learned-order reactivation (Supplementary Figs. S3–S6), exploratory sequenceness–behavior relationships (Supplementary Figs. S7–S8), and an operational candidate-event analysis that complements continuous TDLM sequenceness (Supplementary Fig. S9).

#### Supplementary Methods

##### Behavioral reconstruction and complete-case accounting

Correct-sequence speed was reconstructed from the literal 4–1–3–2–4 keypress logs using the frozen contiguous-motif convention: speed was 10,000 divided by the native-time interval between the first and fifth presses of a correctly completed motif. Trial-mean speed averaged all observed correct motifs within a practice cycle. Rest-linked micro-offline gain (MOG) was defined as the first correct-sequence speed in practice i+1 minus the last correct-sequence speed in practice i, consistent with the brief-rest motor-learning framework [1,2]. Early cumulative MOG summed links 1–11 (cycles 1–12), whereas total cumulative MOG summed all 35 links (cycles 1–36). Neural summaries used the corresponding rest intervals, and negative gains were retained.

Complete early and total behavioral endpoints were available in 61 of 64 participants (No cue 17, Regular 14, Fast 16, Random 14). Two Regular participants and one Random participant lacked the initial correct-sequence endpoint; missing values were not imputed or replaced by partial sums. The early Fast–Regular trial-mean comparison shown in the main manuscript used a two-sided Welch test, with nine auditory-group pair × phase comparisons forming the Bonferroni family. Reconstructed behavioral measures were also compared with published No-cue summaries as a context check because the original scoring code was not part of the data delivery.

#### EEG preprocessing and event alignment

Acquisition procedures are described in the parent study [5]. The present analysis used the 61 scalp EEG channels in the delivered recordings; auxiliary EMG and EOG channels were reserved for sensitivity analyses and did not enter the neural decoders. Continuous EEG was processed with the frozen analysis pipeline using 0.5–100 Hz band-pass filtering, 60-Hz notch filtering, independent-component processing with component classification, and resampling to a 600-Hz working grid. Continuous preprocessing was applied once per recording, whereas feature scaling, class balancing and classifier fitting were restricted to the training data within each cross-validation fold.

Practice keypresses and rest boundaries were aligned from the delivered task markers. Decoder observation windows were required to remain within the relevant practice or rest interval. The analysis used the delivered recording-time coordinates before resampling; detailed preprocessing and auxiliary-channel QC records are retained with the source-data package.

#### Decoder construction and validation details

The primary decoder used 61 scalp EEG channels and two chronological mean-amplitude bins spanning the 150 ms immediately preceding each practice keypress, yielding 122 features. Features were standardized from the training data and classified with participant-specific shrinkage linear discriminant analysis (least-squares solver with automatic covariance shrinkage) using class-balanced training samples. Fivefold validation held out entire practice cycles, so all events from a test cycle were excluded from decoder training. The four-class decoder represented physical fingers 1–4; the five-output full-sequence decoder represented P1–P5 of 4–1–3–2–4 and therefore distinguished the repeated first and final finger 4. P1/P5 discrimination was calculated from held-out five-output posteriors as P5/(P1+P5), without fitting a separate binary classifier.

Three participants had 35 evaluable five-output cycles because cycle-1 position labels were unavailable; all 64 participants remained in the analysis. For the temporal-resolution analysis, the same two-bin feature construction was applied to pre-keypress windows from 25 to 250 ms in 25-ms increments. Final practice-trained participant models were applied to rest on a 5-ms grid without using rest data for fitting.

#### Replay statistics and learned-order reference

Replay sequenceness was quantified using TDLM [3]. Within each intact rest, decoded probability vectors were row-normalized and standardized by state. The frozen first-level model regressed lagged state probabilities on the concurrent probability vector with an intercept using the minimum-norm pseudoinverse (rcond = $10^{-10}$). The resulting lagged association matrix was projected jointly onto Forward (F), Reverse (R), Control (C), self-transition and general-transition templates. We retained F, R and C together with F−R, F−C and R−C. The full-sequence Forward graph was P1→P2→P3→P4→P5, Reverse was P5→P4→P3→P2→P1, and Control was P1→P2→P4→P3→P5. The physical-finger representation used the corresponding 4→1→3→2→4, 4→2→3→1→4 and 4→1→2→3→4 templates. The fixed reference analysis used the pre-keypress 150-ms decoder at a 20-ms transition lag.

Direct lagged covariance was calculated from the same standardized posterior traces by averaging lagged cross-state covariance across the prespecified Forward or Reverse edges and taking their difference. This estimator avoids the first-level predictor-covariance inversion used in TDLM while retaining the same decoded EEG evidence. Learned-order specificity was evaluated against six constrained middle-state assignments that preserved sequence endpoints. Five thousand participant-wise assignments were used to form the conditional reference distributions. Expanded lag-family and joint sensitivity corrections are summarized in Table S3 and the machine-readable source data.

#### Rest support and temporal-resolution analyses

Full-rest analyses retained each nominal 10-s rest as an intact unit. The central-rest sensitivity retained nominal 2–8 s intervals and required the complete classifier observation window and lag pair to remain within the same rest; rests were never concatenated. At W150, lag profiles were evaluated from 5 to 100 ms. The full-cohort temporal-resolution analysis included all 64 participants, all 36 rests, ten decoder windows (25–250 ms), and 50 lags (5–250 ms). For each window, the observed peak was the largest signed group-mean coefficient on the tested grid. Participant-bootstrap distributions characterized peak-location uncertainty. Decoded-state autocorrelation half-width was calculated from the same posterior traces, and W/2 was retained only as a visual guide to the two-bin decoder structure. These analyses directly assess the decoder-dependent temporal structure emphasized in contemporary TDLM methodology [3,4].

#### Replay-behavior statistics

Pooled replay-behavior analyses used the same 61 complete participants at each lag. Ordinary associations were Spearman correlations. Adjusted analyses rank-transformed neural score, gain and baseline performance and residualized neural and behavioral ranks against auditory-condition indicators and ranked initial speed. Nominal ordinary p values used within-group outcome-label permutations; adjusted p values used within-group Freedman-Lane residual permutations. Pointwise 95% intervals used 2,000 participant-bootstrap resamples stratified by cueing group.

Within each decoder representation and adjustment model, the FWER160 family comprised four neural predictors (F, R, C and F−C), 20 lags and two outcomes. The group-specific analysis underlying Supplementary Figure S8 retained Forward, Reverse and Control profiles separately in each cueing group; its FWER960 reference covered four groups × two outcomes × two models × three components × 20 lags.

#### Movement-related sensitivity analyses

Recorded rest-period keypresses were censored with a +/-500-ms guard while preserving complete observation windows and lag pairs. Keypress-censored analyses were available in all 64 participants. An inherited hand-EMG envelope proxy supplied a second movement-related sensitivity analysis with complete 36-rest support in 58 participants (No cue 17, Regular 15, Fast 14, Random 12). Comparisons used matched available participants. Detailed auxiliary-channel availability and preprocessing QC are retained in the source tables.

#### Candidate-event density analyses

Candidate-event analyses provided a complementary event-like summary of the resting decoded-state signal. The replay detector used the five-output pre-keypress W150 decoder, a 150-ms sequence span and a cross-fitted 95th-percentile Control-derived threshold. Local peaks were deduplicated according to the frozen detector rules, and rates were expressed as accepted candidates per valid minute. Forward, Reverse and Control densities were calculated separately from the continuous TDLM coefficients.

Early and total candidate-density associations used the same 11-link and 35-link MOG endpoints where available. Phase-specific analyses used early (links 1–11), intermediate (13–23) and late (25–35) periods, with phase-start performance and cueing condition included where appropriate. Table S6 summarizes the phase-specific between-participant models, while the complete within-participant and nonlinear results are provided as machine-readable Supplementary Data.

#### Source data and reproducibility

Every supplementary figure is accompanied by machine-readable source data and a dedicated plotting script. The accompanying files preserve the complete tested grids and statistical outputs; the summary tables in this document focus on the quantities most directly relevant to interpretation.

Across the supplementary analyses, three themes extend the main findings. First, the behavioral reconstruction and practice-cycle-held-out decoders provide a stable foundation for replay inference. Second, the learned directional signature persists across state representations, estimators and rest supports while its apparent timescale shifts systematically with decoder temporal integration. Third, exploratory sequenceness–gain profiles vary with state representation and cueing context, motivating prospective hypotheses without changing the primary learned-order reactivation result.

##### Supplementary Note 1 | Behavioral reconstruction recapitulates the learning context

The reconstructed behavioral data preserved the characteristic learning trajectory across 36 practice cycles and the early separation between Fast and Regular cueing (Supplementary Fig. S1). The early Fast–Regular difference in trial-mean speed was 0.169 sequences $s^{-1}$ (95% CI 0.087–0.251; Bonferroni-adjusted $p = 0.0022$). Cumulative MOG remained positive across groups, while within-practice micro-online change was negative, reproducing the complementary between- and within-practice dynamics that motivated the replay analysis. The reconstructed No-cue session trial-mean change also closely matched the published summary (0.5035 versus 0.505 sequences $s^{-1}$; Table S4).

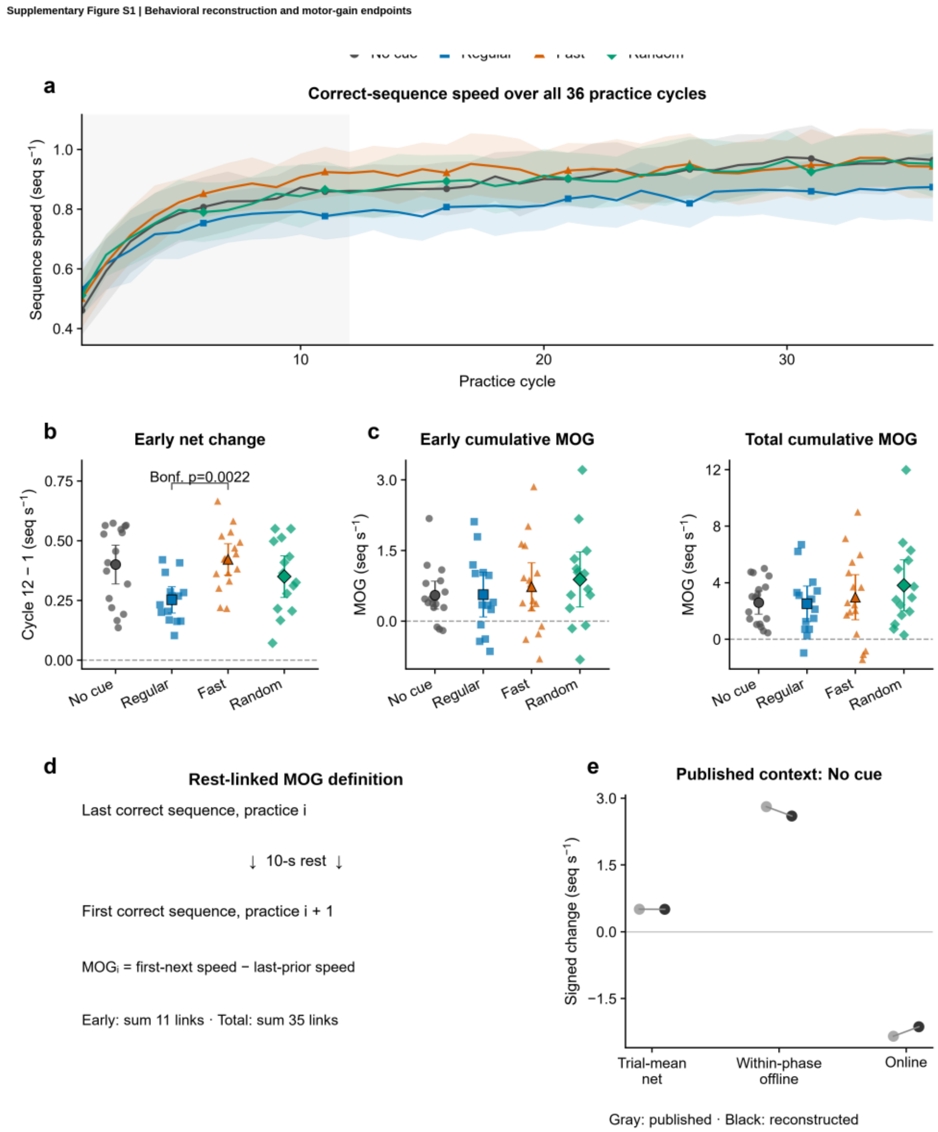


Supplementary Figure S1 | Behavioral reconstruction and motor-gain endpoints. a, Correct-sequence speed across 36 practice cycles. b, Early trial-mean change across cycles 1–12, including the Fast–Regular comparison. c, Early and total cumulative MOG distributions for participants with complete endpoints. d, Definition of rest-linked MOG from the last correct sequence before rest to the first correct sequence after rest. e, Context comparison between published and reconstructed No-cue behavioral summaries. Lines and whiskers show participant-based 95% confidence intervals where applicable. MOG, micro-offline gain.

#### Supplementary Note 2 | Decoder performance is stable across classes and folds

Practice-cycle-held-out decoding was distributed across classes and folds rather than being carried by a single movement or validation partition (Supplementary Fig. S2). Confusion matrices were strongly diagonal for both physical-finger and full-sequence states, and participant balanced accuracy exceeded the corresponding nominal chance level throughout the cohort. The same-finger P1/P5 diagnostic was particularly informative: mean balanced accuracy ranged from 0.766 to 0.808 across groups, showing that the repeated finger 4 carried separable sequence-context information. Fold-level distributions were similar across the five held-out partitions, indicating that no single validation fold dominated decoder performance.

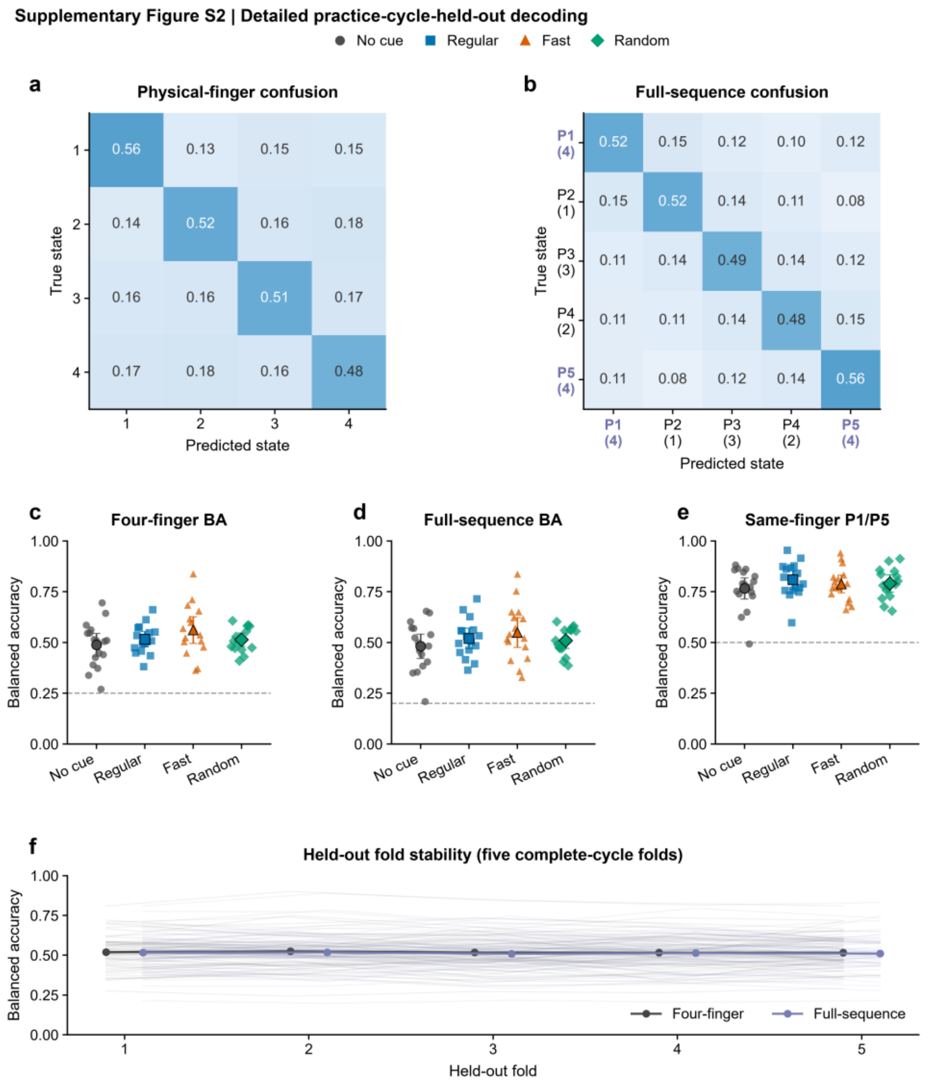


**Supplementary Figure S2 | Detailed practice-cycle-held-out decoding.** a,b, Participant-equally-weighted confusion matrices for four physical fingers and five full-sequence positions. c,d, Participant balanced accuracies with group means and 95% confidence intervals. e, Conditional P1/P5 discrimination from held-out five-output posteriors; P1 and P5 correspond to the same physical finger. f, Fold-level performance across five whole-cycle-held-out partitions. Nominal chance is 0.25, 0.20 and 0.50 for four-finger, full-sequence and P1/P5 decoding, respectively.

### Supplementary Note 3 | Physical-finger representation preserves the learned directional preference

Collapsing the first and final occurrences of finger 4 into a common physical state did not abolish the learned directional signature (Supplementary Fig. S3). At the 20-ms reference lag, four-finger TDLM F−R was positive in each cueing group, and direct covariance showed the same group-level preference. Full-rest and central-rest participant values remained closely aligned. The physical-finger representation therefore provides a complementary view of replay based on action identity, whereas the primary five-output analysis preserves the complete ordinal structure of the practiced sequence.

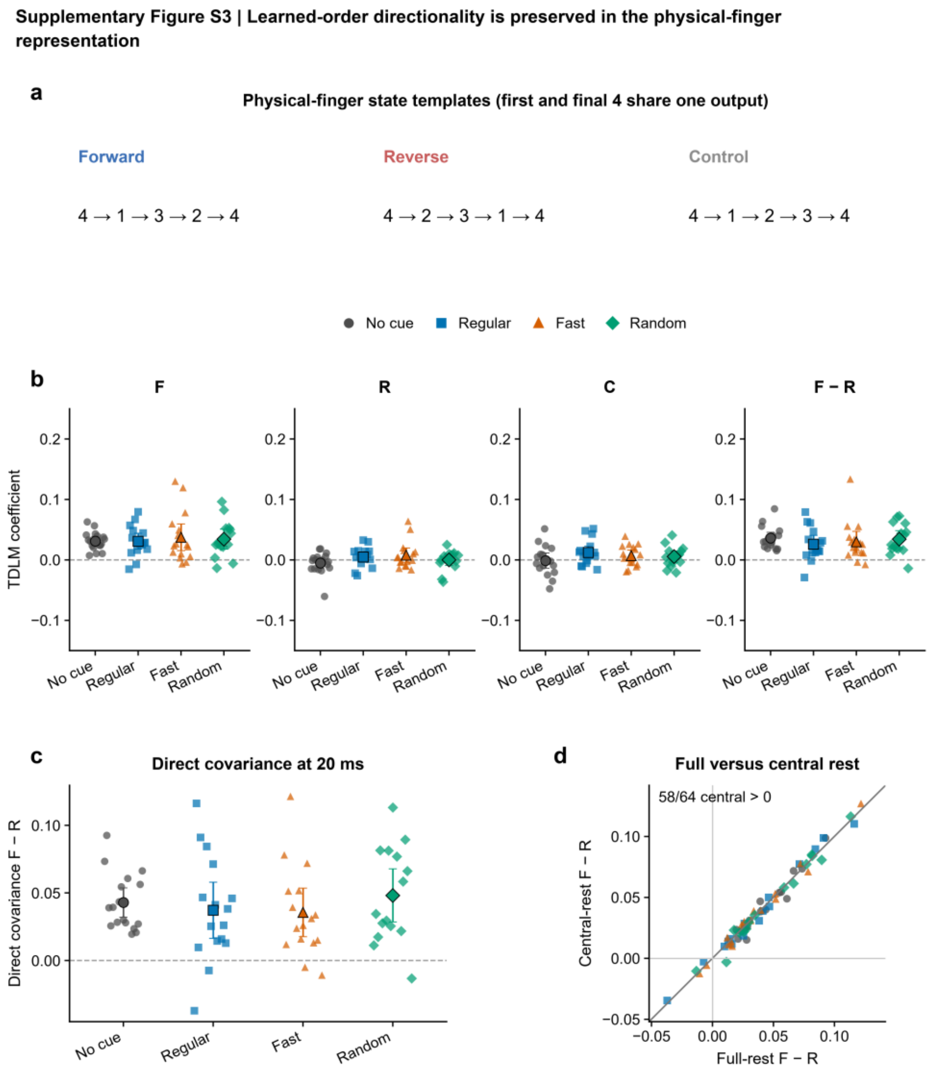


Supplementary Figure S3 | Learned-order directionality is preserved in the physical-finger representation. a, Physical-finger Forward, Reverse and Control transition templates. b, Participant TDLM F, R, C and F−R at the 20-ms reference lag. c, Direct-covariance F−R at the same setting. d, Full-rest versus central 2–8-s direct-covariance F−R with an identity line. The physical-finger decoder collapses the first and final finger 4 and therefore serves as a representation sensitivity to the full-sequence analysis.

#### Supplementary Note 4 | Learned-order directionality across lag, estimator and rest support

At the fixed full-sequence setting, direct-covariance F−R was positive in 61 of 64 participants, and the practiced template ranked first among the six constrained orders in 52 of 64 (Table S2). The directional replay signature was not confined to the historical 20-ms reference point. Across 5–100 ms, full-sequence F−R increased toward a broad maximum in each cueing group, and the temporal profile was closely reproduced by direct covariance (Supplementary Fig. S4a–c). Full-rest and central-rest profiles were also similar (Supplementary Fig. S4d), indicating that the learned-order organization was distributed across the resting interval rather than restricted to its immediate boundary. At each group peak, the practiced template ranked first among the six permitted assignments, with peak F−R values of 0.143–0.166 at 85–90 ms (Table S3).

**Supplementary Figure S4 | Learned-order directionality persists across lag, estimator and rest support**

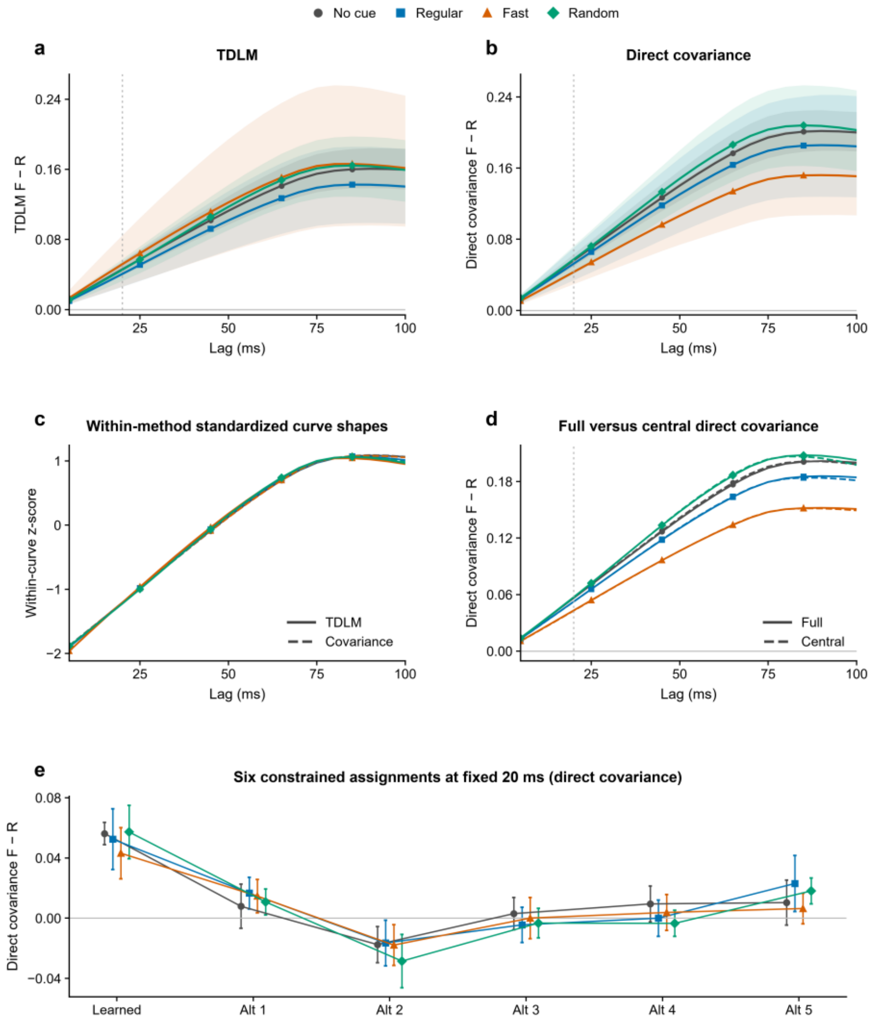


Supplementary Figure S4 | Learned-order directionality persists across lag, estimator and rest support. a,b, Full-sequence F−R across 5–100-ms lags for TDLM and direct lagged covariance in all 64 participants. c, Standardized within-method curves compare temporal shape across estimators. d, Full-rest and central 2–8-s direct-covariance profiles. e, Practiced and constrained alternative sequence assignments at the 20-ms reference setting. Shading denotes pointwise participant-bootstrap 95% confidence intervals where shown.

#### Supplementary Note 5 | Movement-related censoring leaves learned-order directionality largely unchanged

Recorded keypresses during nominal rest were concentrated near rest boundaries, and applying the predefined keypress guard produced minimal changes in full-sequence F−R (Supplementary Fig. S5a,b). The independent EMG-proxy sensitivity, available in 58 participants with complete support, produced similarly small paired changes (Supplementary Fig. S5c,d). Across both checks, participant estimates remained close to the identity line, showing that the learned directional signature is stable to the recorded movement-related activity captured by these controls.

Supplementary Figure S5 | Learned-order directionality is stable to movement-related censoring

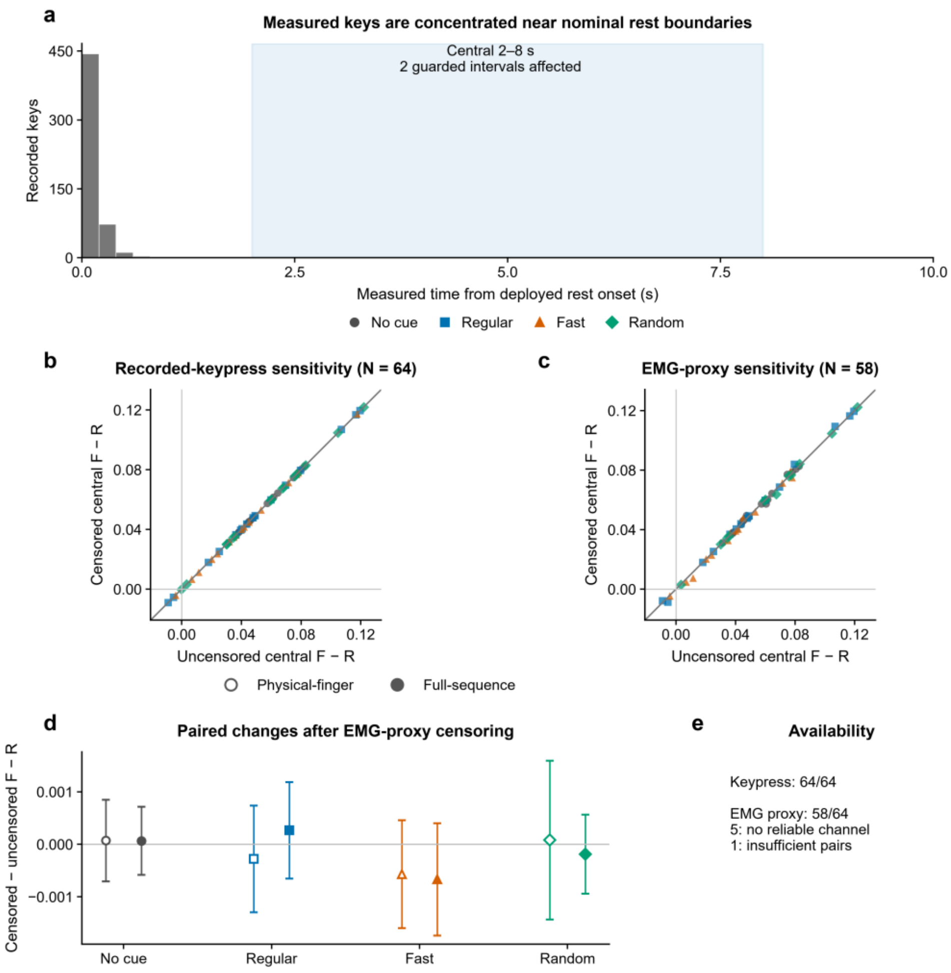


Supplementary Figure S5 | Learned-order directionality is stable to movement-related censoring. a, Timing of recorded keypresses relative to nominal rest. b, Paired full-sequence direct-covariance F−R before and after keypress censoring in all 64 participants. c, Corresponding EMG-proxy sensitivity in 58 participants with complete 36-rest coverage. d, Group-level paired changes for physical-finger and full-sequence representations. e, EMG-proxy availability by cueing group. Identity lines denote unchanged values.

#### Supplementary Note 6 | Decoder temporal integration shapes the measured sequenceness timescale

The full-cohort window sweep revealed a highly ordered relationship between decoder duration and the apparent Forward sequenceness peak (Supplementary Fig. S6). In the full-sequence representation, the Forward maximum moved from 15 ms at W25 to 130 ms at W250, passing through 80 ms at the primary W150 setting. The physical-finger representation showed a closely related trajectory. F−C broadly followed the same ridge, whereas F−R could exhibit competing late maxima at short windows because its value depends on both Forward and Reverse components. Decoded-state autocorrelation broadened in parallel with decoder duration, reinforcing the interpretation that the measured temporal profile is constrained by decoder integration.

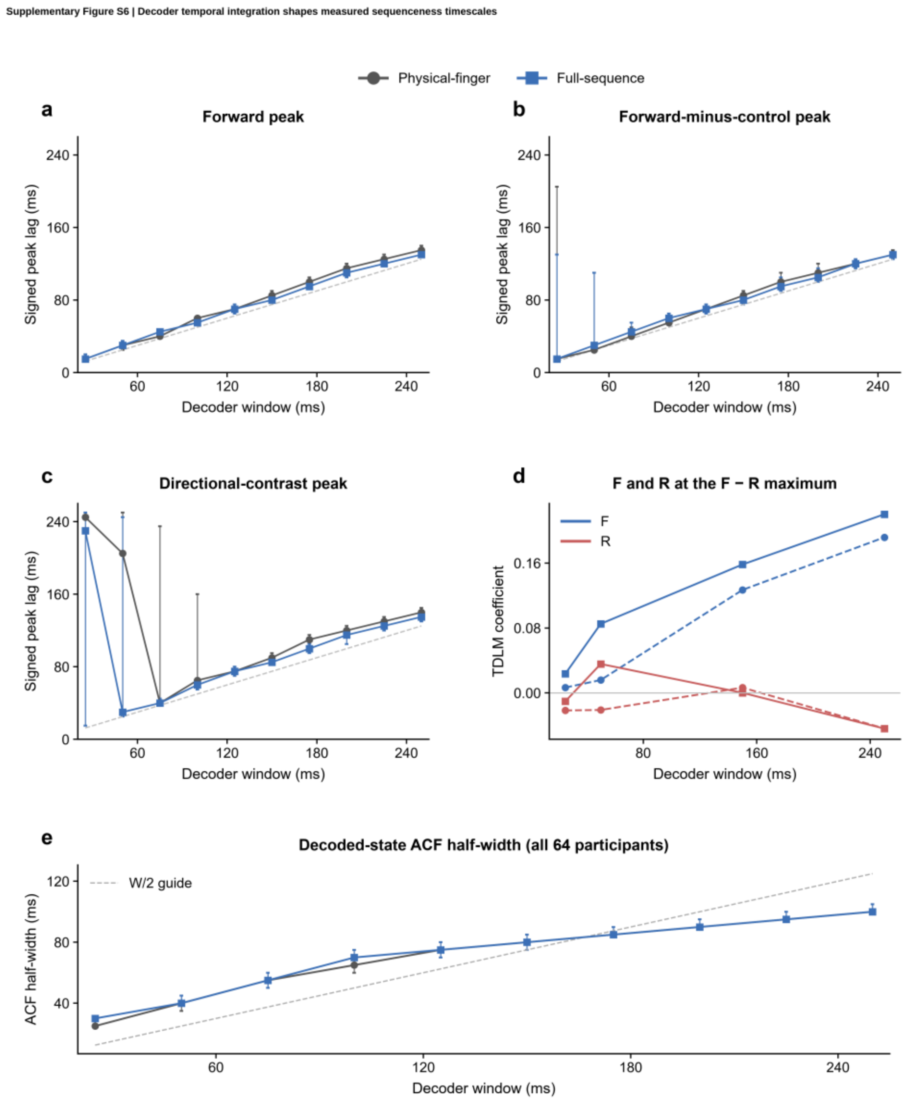


Supplementary Figure S6 | Decoder temporal integration shapes measured sequenceness timescales. a–c, Forward F, F−C and F−R peak lags across 25–250-ms decoder windows in both representations, using all 64 participants and the 5–250-ms lag grid. d, Forward and Reverse values evaluated at the F−R maximum for representative windows. e, Decoded-state autocorrelation half-width across decoder windows. The W/2 line is a visual guide to the two-bin observation model. Whiskers show participant-bootstrap peak-location uncertainty.

#### Supplementary Note 7 | Full-sequence sequenceness shows a complementary behavioral profile

Behavioral coupling depended on how the motor sequence was represented. In the full-sequence decoder, the largest ordinary early association was Forward sequenceness at 35 ms ($\rho = 0.280$, nominal $p = 0.030$; Supplementary Fig. S7; Table S5). Adjusted relationships were smaller than those in the physical-finger representation highlighted in main Figure 5, and no selected full-sequence association survived the FWER160 reference. The two state spaces therefore provide complementary descriptions of the resting decoded signal rather than independent confirmatory behavioral effects. Forward-versus-Control correlation differences and the correlation of F−C with gain are shown separately because they address distinct estimands.

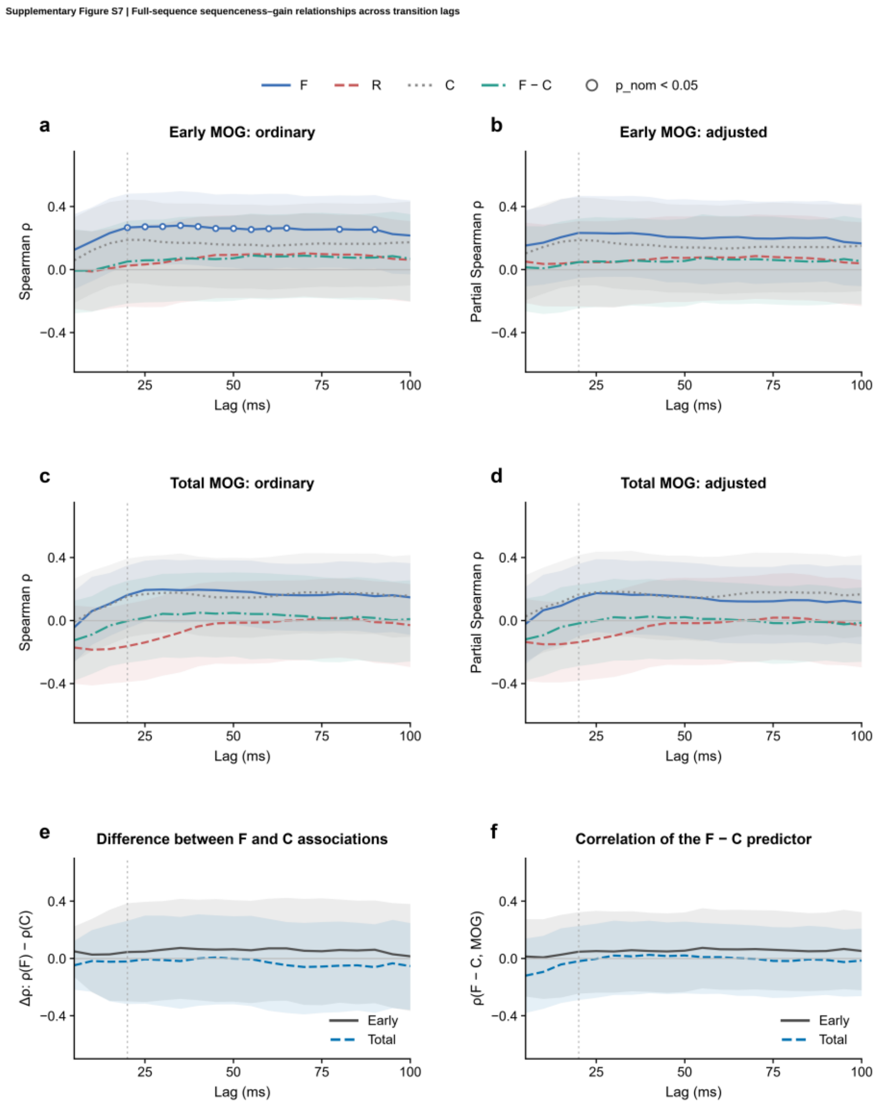


Supplementary Figure S7 | Full-sequence sequenceness–gain relationships across transition lags. a–d, Ordinary and adjusted relationships between full-sequence TDLM coefficients and early or total cumulative MOG in the same 61 participants. Shading denotes pointwise participant-bootstrap 95% confidence intervals; open symbols indicate nominal permutation $p < 0.05$. e, Paired-bootstrap difference between Forward and Control correlations. f, Correlation of the F−C predictor with gain. The historical 20-ms lag is shown for reference.

#### Supplementary Note 8 | Exploratory group-specific sequenceness–gain profiles

Cueing groups showed descriptively different sequenceness–gain profiles when examined separately (Supplementary Fig. S8). In the Fast group ($n = 16$), the retrospectively selected 5-ms Forward association with total cumulative MOG was positive (ordinary Spearman $\rho = 0.747$; nominal $p = 0.0008$), but the group-specific FWER960 reference was not significant ($p = 0.165$).

Because the lag was selected retrospectively and no formal between-group interaction was established, this pattern is presented only as a descriptive subgroup observation and a target for prospective testing.

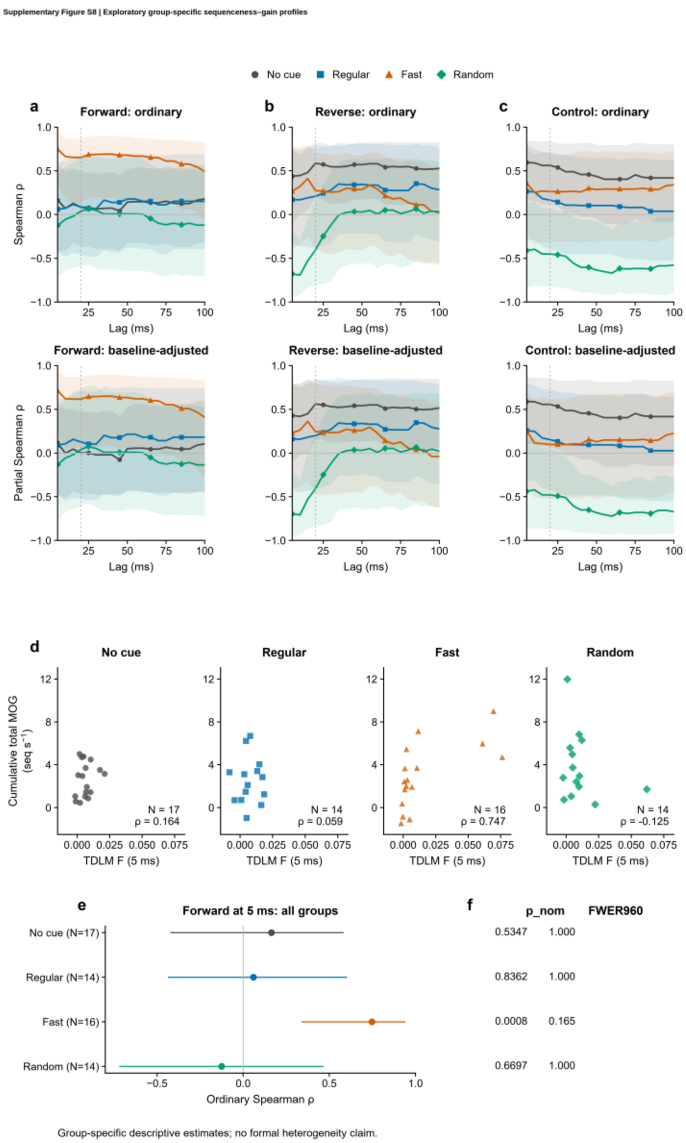


Supplementary Figure S8 | Exploratory group-specific sequenceness–gain relationships. a–c, Ordinary and baseline-adjusted Spearman relationships between Forward, Reverse and Control four-finger TDLM coefficients and total cumulative MOG across 5–100-ms lags, shown separately for each cueing condition. d, Participant-level Forward observations at the retrospectively selected 5-ms lag used in main Figure 5. e,f, Group-specific Forward correlations with 95% confidence intervals and the saved nominal and family-wise references. Group Ns are No cue 17, Regular 14, Fast 16 and Random 14. The Fast 5-ms association is descriptive and does not survive the FWER960 reference; no formal between-group interaction is claimed.

#### Supplementary Note 9 | Operational candidate-event density provides a complementary summary

The candidate detector yielded frequent Forward, Reverse and Control candidates across all cueing groups (Supplementary Fig. S9a), providing an event-like summary of the decoded resting signal that is analytically distinct from continuous TDLM. Forward candidate density was numerically higher than Reverse density, whereas Control density was similar to Forward in several groups. Candidate-density relationships with early and total MOG were modest in the pooled analyses (Supplementary Fig. S9b,c), indicating that continuous sequenceness and thresholded candidate density emphasize different features of the resting decoded-state activity.

**Supplementary Figure S9 | Operational candidate-event density and motor-gain relationships**

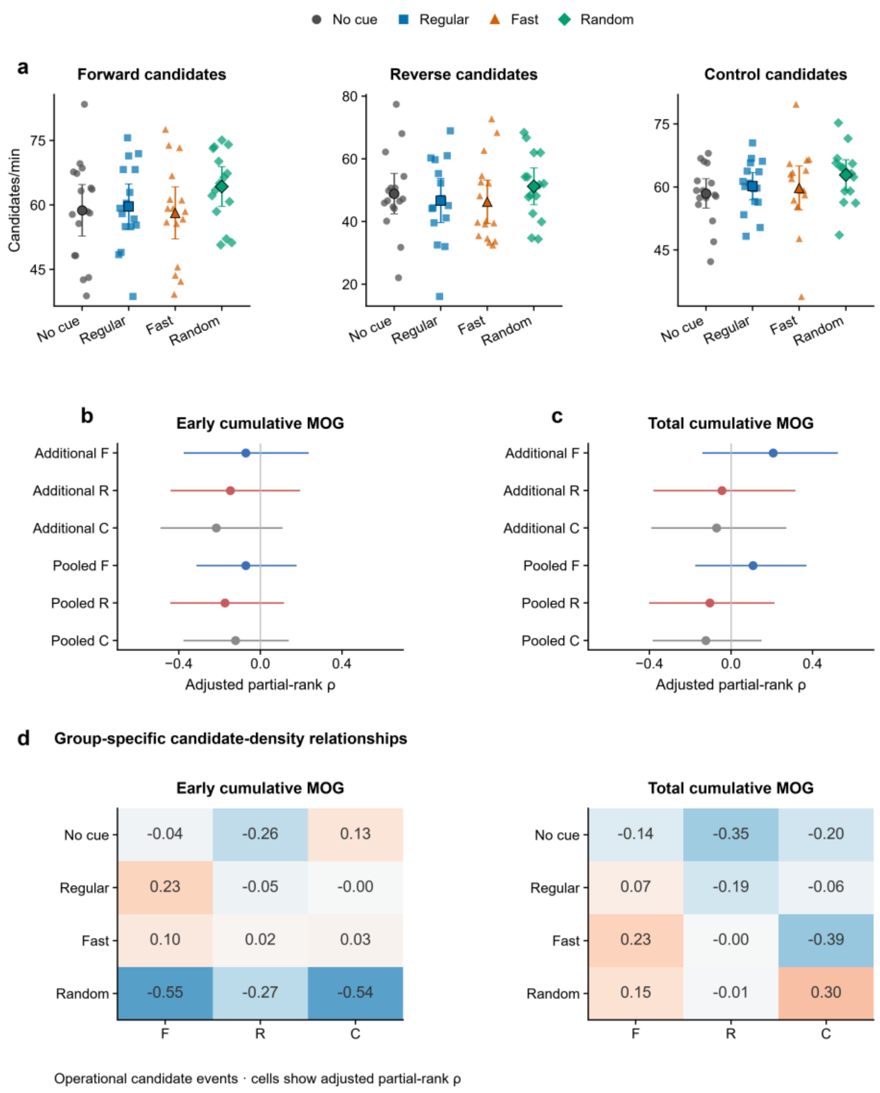


Supplementary Figure S9 | Operational candidate-event density and motor-gain relationships. a, Forward, Reverse and Control operational candidate-event densities across rests 1–35. b,c, Group- and baseline-adjusted candidate-density relationships with early and total cumulative MOG in the additional auditory cohort and pooled cohort. d, Group-specific adjusted relationships for Forward, Reverse and Control under both outcomes. Candidate density is an operational detector output and is analytically distinct from continuous TDLM sequenceness; phase-specific analyses are summarized in Table S6 and the accompanying Supplementary Data.

Phase-specific models included several nominal positive coefficients, including Fast intermediate-phase Forward density (β = 0.773) and Forward−Control (β = 1.401), and late-phase effects in the additional auditory cohort (Table S6). None survived the Holm150 correction. These estimates are therefore treated as exploratory targets for prospective analyses rather than evidence for phase- or cue-specific candidate-event effects.

#### Supplementary Note 10 | Synthesis across supplementary analyses

Together, the supplementary analyses support the central neural result from multiple directions: practice decoding is stable across classes and folds; learned-order directionality is retained in the physical-finger representation, reproduced with direct covariance and preserved under rest-boundary and movement-related sensitivities; and decoder-window analyses show that the apparent sequenceness timescale depends on temporal integration. The behavioral analyses remain exploratory and representation dependent, and the group-specific and candidate-density findings do not survive their declared family-wise corrections. The strongest evidence therefore remains the reproducible learned-order structure of decoded scalp EEG during wakeful rest.

### Supplementary Tables

##### Supplementary Table S1 | Participant groups and practice-cycle-held-out decoder performance

| Condition | N | Four-finger BA | Full-sequence BA | P1/P5 BA |
|---|---|---|---|---|
| No cue | 17 | 0.489 | 0.481 | 0.766 |
| Regular | 16 | 0.515 | 0.520 | 0.808 |
| Fast | 16 | 0.561 | 0.549 | 0.788 |
| Random | 15 | 0.514 | 0.509 | 0.791 |

*Nominal chance is 0.25, 0.20 and 0.50, respectively. Values are participant-level group means.*

Supplementary Table S2 | Fixed-setting full-sequence sequenceness summary at 20 ms

| Condition | N | TDLM F−R | Direct covariance F−R | Positive DC F−R | Learned rank 1/6 |
|---|---|---|---|---|---|
| No cue | 17 | 0.0459 | 0.0562 | 17/17 | 16/17 |
| Regular | 16 | 0.0406 | 0.0525 | 14/16 | 12/16 |
| Fast | 16 | 0.0517 | 0.0431 | 15/16 | 11/16 |
| Random | 15 | 0.0453 | 0.0572 | 15/15 | 13/15 |

*Five-output/full-sequence decoder, W150, all 36 rests. TDLM and direct covariance use different coefficient scales.*

##### Supplementary Table S3 | Lag-resolved conditional learned-order reference

| Condition | N | Peak lag (ms) | Peak F-R | FWER80 p | Joint FWER640 p |
|---|---|---|---|---|---|
| No cue | 17 | 90 | 0.161 | 0.0002 | 0.0002 |
| Regular | 16 | 85 | 0.143 | 0.0002 | 0.0008 |
| Fast | 16 | 85 | 0.166 | 0.0310 | 0.0832 |
| Random | 15 | 85 | 0.164 | 0.0002 | 0.0002 |

Full-sequence TDLM, W150, full rest. The table summarizes the saved lag-family reference used for the expanded 5–100-ms analysis.

**Supplementary Table S4 | Context check for reconstructed No-cue behavioral summaries**

| Measure | N | Published | Reconstructed | Difference |
|---|---|---|---|---|
| Session trial-mean gain | 17 | 0.505 | 0.504 | -0.001 |
| Within-phase micro-offline | 17 | 2.809 | 2.600 | -0.209 |
| Within-phase micro-online | 17 | -2.348 | -2.136 | +0.212 |

The primary replay–behavior endpoints are the 11-link early and 35-link total MOG measures; this table provides context for the reconstruction. Values are expressed in sequences $s^{-1}$.

**Supplementary Table S5 | Selected full-sequence sequenceness–gain associations**

| Outcome | Model | Measure | Lag (ms) | ρ | Nominal p | FWER160 p |
|---|---|---|---|---|---|---|
| Early | Ordinary | F | 35 | 0.280 | 0.030 | 0.363 |
| Early | Ordinary | R | 70 | 0.103 | 0.440 | 0.999 |
| Early | Adjusted | F | 20 | 0.232 | 0.080 | 0.672 |
| Early | Adjusted | R | 70 | 0.086 | 0.536 | 1.000 |
| Total | Ordinary | F | 30 | 0.197 | 0.139 | 0.813 |
| Total | Adjusted | F | 25 | 0.173 | 0.210 | 0.933 |
| Total | Adjusted | R | 80 | 0.018 | 0.895 | 1.000 |

Rows show the largest signed correlation for the indicated full-sequence predictor/model across 5–100 ms. Complete lag profiles are shown in Supplementary Figure S7 and provided as source data.

**Supplementary Table S6 | Selected phase-specific candidate-density models**

| Cohort | Phase | Predictor | N | Effect | 95% CI | Nominal p | Holm150 p |
|---|---|---|---|---|---|---|---|
| Fast | Intermediate | Forward unique | 16 | 0.773 | [0.242, 1.385] | 0.0282 | 1.000 |
| Fast | Intermediate | Forward-Control | 16 | 1.401 | [0.410, 2.641] | 0.0356 | 1.000 |
| Additional auditory cohort | Late | Forward unique | 47 | 0.502 | [0.109, 0.809] | 0.0106 | 1.000 |
| Additional auditory cohort | Late | Forward-Control | 47 | 0.862 | [0.150, 1.339] | 0.0098 | 1.000 |
| Pooled cohort | Late | Forward unique | 64 | 0.313 | [-0.046, 0.642] | 0.0984 | 1.000 |
| Pooled cohort | Late | Forward-Control | 64 | 0.469 | [-0.131, 1.009] | 0.1412 | 1.000 |

*Effects are standardized rank-model coefficients from the frozen phase-specific analysis. The complete between-participant, within-participant and nonlinear tables are supplied as machine-readable Supplementary Data.*

#### Supplementary Data

Complete source-data CSVs for Supplementary Figures S1–S9, the phase-specific candidate-density tables and the plotting scripts accompany this document. The summary tables are intentionally focused; machine-readable files preserve the complete tested grids and reported statistical outputs.

### Supplementary References


1. Bönstrup, M. et al. A rapid form of offline consolidation in skill learning. Current Biology 29, 1346–1351.e4 (2019). https://doi.org/10.1016/j.cub.2019.02.049
2. Das, A., Karagiorgis, A., Diedrichsen, J., Stenner, M.-P. & Azañón, E. Micro-offline gains do not reflect offline learning during early motor skill acquisition in humans. Proceedings of the National Academy of Sciences USA 122, e2509233122 (2025). https://doi.org/10.1073/pnas.2509233122
3. Liu, Y. et al. Temporally delayed linear modelling (TDLM) measures replay in both animals and humans. eLife 10, e66917 (2021). https://doi.org/10.7554/eLife.66917
4. Kern, S. et al. Challenges in replay detection by TDLM in post-encoding resting state. eLife 14, RP108023 (2026). https://doi.org/10.7554/eLife.108023
5. Kawasoe, R. et al. Wakeful targeted memory reactivation during short rest periods modulates early motor learning. npj Science of Learning 11, 23 (2026). https://doi.org/10.1038/s41539-026-00407-9
